# Coupling of 12 chromosomal inversions maintains a strong barrier to gene flow between ecotypes

**DOI:** 10.1101/2023.09.18.558209

**Authors:** Alan Le Moan, Sean Stankowski, Marina Rafajlovic, Olga Ortega-Martinez, Rui Faria, Roger Butlin, Kerstin Johannesson

## Abstract

Chromosomal rearrangements lead to the coupling of reproductive barriers, but whether and how they contribute to completion of speciation remains unclear. Marine snails of the genus *Littorina* repeatedly form hybrid zones between taxa segregating for inversion arrangements, providing opportunities to study this question. Here, we analysed two adjacent transects across hybrid zones between large and dwarf ecotypes of *Littorina fabalis* covering wave exposure gradients in a Swedish island. Applying whole-genome sequences we found 12 putative inversions reaching near differential fixation between the opposite ends of each transect, and being in strong linkage disequilibrium. These inversions cover 20% of the genome and carry 93% of divergent SNPs. Bimodal hybrid zones in both transects indicate that the two ecotypes of *Littorina fabalis* maintain their genetic and phenotypic integrity following contact due to strong coupling between inversion clines that strengthened the reproductive barrier. The bimodality resulting from the linked inversions extends into collinear parts of the genome, suggesting a genome-wide coupling. Demographic inference suggests that the coupling built up during a period of allopatry, and has been maintained for more than 1K generations after secondary contact. Overall, this study shows that the coupling of multiple chromosomal inversions contributes to strong reproductive isolation. Importantly, two of the inversions overlap with inverted genomic regions associated with ecotype differences in a closely-related species (*L. saxatilis*), suggesting the same regions, with similar structural variants, repeatedly contribute to ecotype evolution in distinct species.

## Introduction

Understanding reproductive isolation (RI) from genome-wide polymorphism data has become a key goal in speciation research^1^. RI evolves in response to divergent natural and sexual selection, and through mutation order effects, both of which lead to the accumulation of barriers to gene flow along the genome^2,3^. The strength of the barrier experienced by a locus depends on its proximity to a causal barrier locus, the fitness effects of the barrier loci, the genome-wide distribution of these barriers, and their interaction in recombinant genomes^4^. These recombinant genomes can be found in hybrid zones, which represent natural laboratories in which to study RI^5^. When RI is weak, a high level of admixture between the parental lineages is expected in the centre of the hybrid zone, which leads to the formation of a unimodal hybrid zone^6^. In contrast, when RI is very strong, hybrids are rare and parental genotypes coexist in the centre of the hybrid zone, which leads to a bimodal hybrid zone^7,8^. Studying the formation of bimodal hybrid zones can provide an important step towards understanding the completion of speciation^9^.

The formation of bimodal hybrid zone requires coupling of barrier loci, that is the build-up of strong linkage disequilibrium (LD) between pre- and post-zygotic barriers to gene flow^10^. This coupling can easily occur during an allopatric phase of divergence prior to secondary contact and hybrid zone formation, or, through reinforcement when divergence occurs in the face of gene flow^11^. In addition, barriers of different geographical origin can overlap in space due to the attraction of allelic clines to the same ecotone, a process coined “spatial coupling”^12,13^. Coupling can also be facilitated when individual barrier loci cluster within genomic regions where recombination is suppressed, for example, within chromosomal inversions^14^. This genomic coupling effectively combines many individual barrier loci into one large effect locus^15^, and also extends local barrier effects across larger segments of chromosome^16,17^. By maintaining LD among barrier loci in the presence of gene flow, such rearrangements facilitate the spread of locally adaptive alleles^17^, promote the accumulation of genetic incompatibilities^18^, and maintain local adaptation. For instance, chromosomal inversions are often associated with the evolution of partially-isolated ecotypes (e.g. deer mouse^19^, Atlantic cod^20^, marine snails^21^, sunflower^22^). However, the role of inversions in generating strong isolation remains unclear.

Marine snails of the genus *Littorina* are emerging models for studying the role of chromosomal inversions in speciation^21,23,24^. *Littorina* includes several species that form ecotypes that differ in multiple phenotypic traits (including shell size, shell morphology, and behaviour). The ecotypes evolve repeatedly across similar types of meter-scale environmental gradients^25^. In *L. saxatilis*, the parallel evolution of ‘wave’ and ‘crab’ ecotypes involves more than 10 chromosomal inversions^21,26^. Many of these show signatures of divergent selection, and contribute to phenotypic differences between the ecotypes^24,27–29^. Despite steep frequency clines across ecotype hybrid zones, the coupling of the different inversions is weak In *L. saxatilis*, and hybrid zones appear unimodal^24,29,30^. In addition, several inversions remain polymorphic within one or both ecotypes, and they only explain roughly half of the phenotypic variation among snails^27^. Therefore, despite much of the genetic divergence underlying local adaptation being associated with the inversions, it remains unclear whether the establishment of these chromosomal rearrangements also generates strong RI.

A closely-related species, *Littorina fabalis*, living adjacent to *L. saxatilis* but in the seaweeds, forms a dwarf and a large ecotype associated to wave exposure gradients on multiple European shores^31,32^. In contrast to the unimodal hybrid zones of *L. saxatilis, L. fabalis* shows some evidence of bimodality in ecotype contact zones^33^. Sharp allele frequency clines separate the ecotypes at one out of four highly polymorphic allozyme loci^33,34^. In Sweden, this allozyme locus shows a deficiency of heterozygotes, and the size distribution is bimodal in the hybrid zone, both suggesting that ecotypes are partly reproductively isolated^33^. Recently, we found that the allozyme gene is located inside a large chromosomal inversion^23^, and preliminary data suggested the presence of additional inversions. Here, we study the genomic landscape of differentiation between the two *L. fabalis* ecotypes across the whole genome using 295 snails along parallel transects from two adjacent hybrid zones between the large and the dwarf ecotypes of *L. fabalis*, located on either side of a small island on the Swedish west coast, Figure 1a). Using low coverage Whole Genome Sequencing (lcWGS), we found that most genetic differences between the ecotypes were localised inside 12 large putative chromosomal inversions. We map the inversion arrangement frequencies across the hybrid zones to assess their barrier effects, and show that the coupling between the differentially fixed inversions results in strong reproductive isolation associated with the formation of bimodal hybrid zones. We furthermore explore the evolutionary origin of the two ecotypes of *L. fabalis*, and provide a comparative genomics analysis of ecotypic divergence with the related *L. saxatilis*.

**Figure 1:**
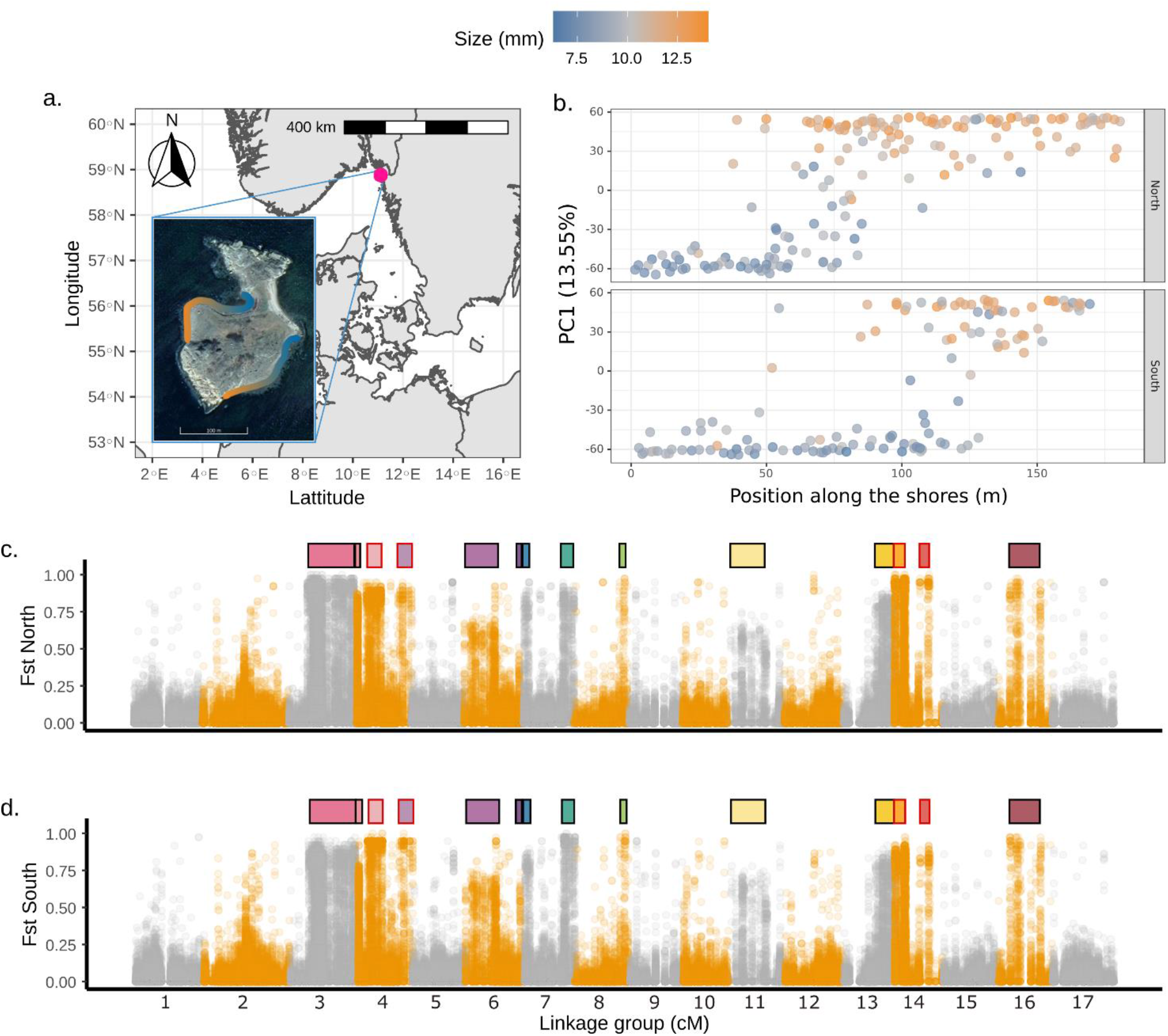
Population structure analyses showing: a. map of the sampling locations; b. the PC1 score of a PCA performed on the 295 snails genotyped at 110k SNPs (filtered set, see Methods) with MAF of >5% that are placed on the linkage map. The individual PC1 scores were plotted against the positions of the snails on the northern (top) and southern (bottom) transects; c. and d. Manhattan plots of pairwise *F*_ST_ calculated between 20 snails sampled from the opposite ends of each transect (top: northern and bottom: southern transect). In a. and b., the colour gradient represents the size of individual snails (following the legend above the graph). In c. and d., the orange and grey show the limits of different linkage groups on which the contigs were ordered by linkage map position, and the horizontal bars on top of the plot show the positions of the 14 blocks (one colour per block) corresponding to 12 putative chromosomal inversions (after grouping the two pairs of blocks with LD of 1 from LG4 and LG14 encircled in red, see text, Figure S2).

## Results

### Population structure and the landscape of differentiation

The two ecotypes of *L. fabalis* were clearly genetically distinct on the first axis of a PCA analysis, which explained a large part of the total variation (PC1 = 13.55%, PC2 < 1%, Figure S1) and showed very similar clines in the two transects (Figure 1b). The average *F*_ST_ between ecotypes, calculated between 20 snails collected from each transect end, was 0.090 (0.092 and 0.088 across the northern and southern shore transects, respectively). Rather than being randomly distributed across the genome, we found that highly differentiated loci formed clusters on 9 of the 17 chromosomes when mapping the sequences to the *L. saxatilis* reference genome (Figure 1c,d).

### The signatures of chromosomal rearrangements

The clusters of high differentiation across the genome might be caused by large chromosomal inversions. Therefore, we looked for signatures commonly associated with inversions^21,35,36^. First, we found strong LD among SNPs located several cM away, revealing 14 LD blocks ranging from 1 to 33Mbp (figure S2, table S1). These blocks were characterized by plateaus of similarly high LD typical of large inversions (Figure S3). The average LD was much stronger within blocks than the average LD found outside (r^2^ = 0.387 and 0.006, respectively, Table S1 and Figure S4). Second, Local PCAs performed on the SNPs within each block revealed three distinct clusters of individuals on PC1 (Figure S5 to S7). Observed heterozygosity was approximately twice as high in the central cluster compared with those on either side (Figure S8 to S10, and Table S2). Moreover, hundreds of SNPs were differentially fixed between the two distant clusters (Table S1). All of these observed patterns are hallmarks of chromosomal inversions. Third, as shown earlier, the largest block located on LG3 shows a suspension bridge pattern^23^, providing further support for this block being an inversion. In some local PCAs (e.g., for LG14_inv2 and to some extent for LG7_inv, Figure S6), more complex patterns of clustering were observed, suggesting the presence of multiple overlapping polymorphic inversions. In addition, LD suggests that separate LD blocks on LG4 and LG14 are probably part of the same inversion, but have been broken up due to differences in synteny between the reference used (*L. saxatilis*) and *L. fabalis* (Figure S11). One additional island of high differentiation was visible near the centre of LG2 (Figure 1). The genomic signatures were different from those of the putative inversions (Figure S2 to S5), and this may be a region of low recombination associated with the centromere^37,26^.

### Consequences of chromosomal rearrangements for population structure

The 12 putative chromosomal inversions (after combining the two blocks on LG4 and the two on LG14) covered around 20% of the reference genome (24% of the linkage map positions and 20% of the base pairs analysed here). The differentiation between ecotypes inside these putative inversions was an order of magnitude higher (mean *F*_ST_ ∼ 0.3) than outside of them (mean *F*_ST_ ∼ 0.03). In addition, they carried 94% of the SNPs with *F*_ST_ values above the 95% upper quantile of differentiation (i.e. *F*_ST_ ≥ 0.61 and 0.59 in the northern and southern transect, respectively), and 92% of these SNPs were shared among the two transects. LD between arrangements located on different linkage groups was also very high (0.35> LD_between_<1, Figure S11), although some inversions had substantially lower LD than others (e.g. inversions on LG6 and LG11). These results showed that 12 putative inversions carry most of the divergence between the two ecotypes in *L. fabalis*. Nevertheless, the ecotypes remained distinct after removing SNPs within the putative inversions (*F*_ST_∼0.03), as illustrated by the first axis of a PCA performed on the collinear part of the genome (only capturing 2% of the variation; Figure S1b).

### Arrangement and allelic clines

Sampling was conducted over continuous transects allowing exploration of the strength of the barrier effect provided by the putative inversions from the steepness of the clines. We inferred arrangement-frequency clines at the 12 putative inversions, using information from clusters inferred in a local DAPC to obtain the karyotypes of each snail (Figure S7). All rearrangement clines showed similar trends with cline centres in close proximity to the phenotypic shift and with different arrangement frequencies reaching near fixation at transect ends in both shores, with the exception of arrangements on LG6 and LG11 that remained polymorphic in the dwarf ecotype (Figure 2d, Table S6). In addition, 94% of the clinal SNPs were found within the inversions and showed similar trends to the rearrangement clines (Figure S18). Using an estimate of dispersal of σ = 8.27 (± 2.01 SE) directly inferred from the distances between 19 related individuals sampled along the transects (10 half-sib pairs and 1 full sib pair, see details MatSup p28), we estimated that a coefficient of selection ranging from 0.02 to 0.09 was required to maintain the arrangement clines. All the differentially fixed rearrangements showed strong and significant deficiencies of heterokaryotypes near the centre of the clines (inferred directly from the cline fitting, Table S6). Nevertheless, all inversions were found as heterozygotes in the centre of the hybrid zone. By representing the hybrid index against the heterozygosity calculated from the inversion karyotypes, we found that inversions were heterozygous in F1 and backcross individuals between the two ecotypes (i.e., individuals located on the top and along the sides of a triangle plot; Figure 2b), but that recombinant individuals from the two genetic backgrounds were largely missing (F2 or latter generations of recombinant hybrids, located in the centre of the triangle plot).

**Figure 2:**
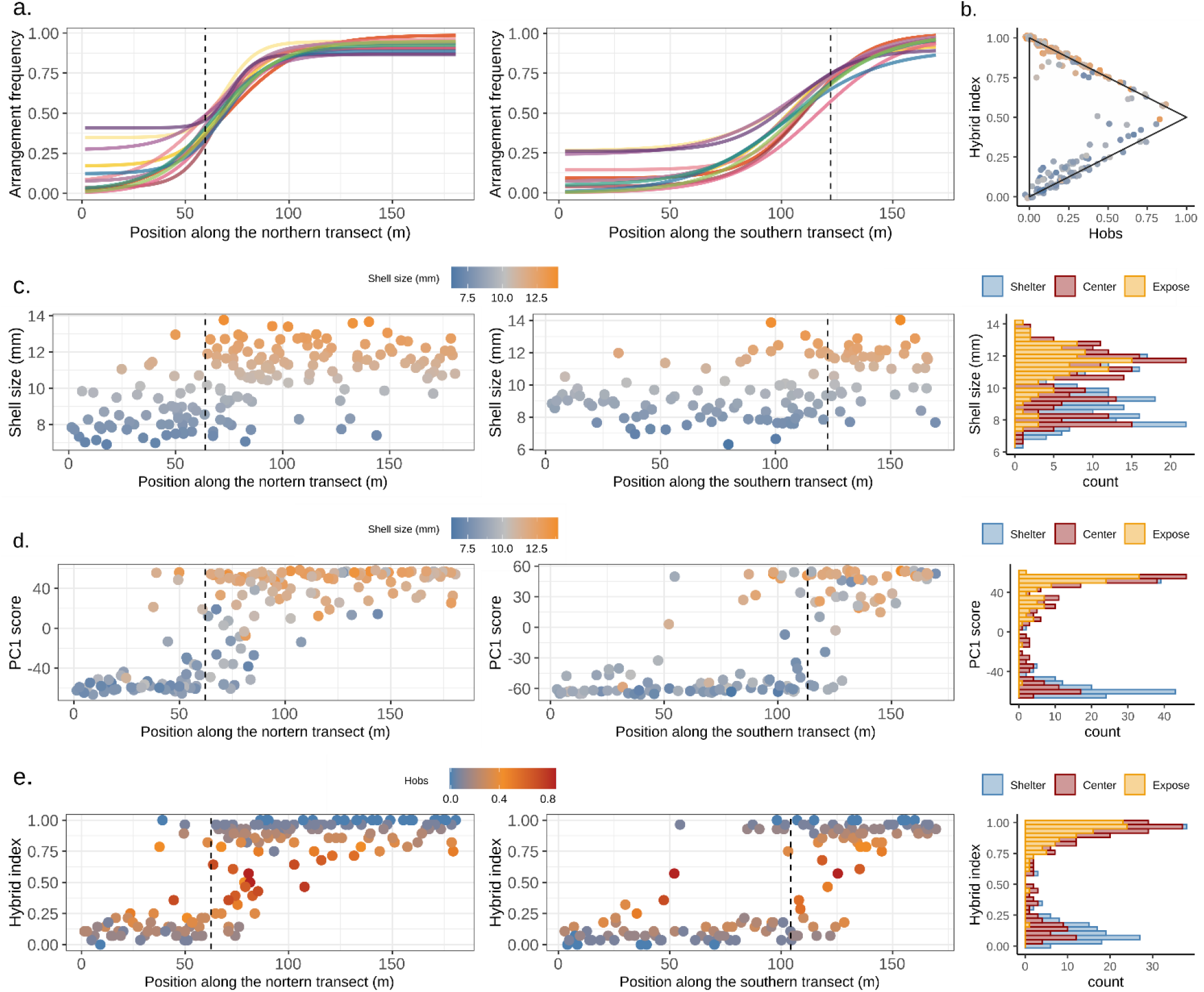
Cline analyses for the northern (left) and southern transect (middle). a. shows frequency clines inferred from the 12 putative chromosomal rearrangements, which are coloured according to the colour rectangles used in Fig. 1c,d. The dotted vertical line in each of the top graphs show the centre of the size phenotypic cline. Graph b. shows the triangle plot for hybrid detection with F1 hybrids expected at the “top right” of the triangle (both transects pooled). The graph b., c., and d., respectively show the shell size variation, the PC1 score variation (as in Figure 1b), and, c. the hybrid index, here calculated from the chromosomal rearrangement karyotypes. The dashed line shows the centre of the cline (Table S3-S5). All the histograms (b. c. and d. on the right, both transects pooled) separated individuals from the middle of the hybrid zone in red (centre of the cline ± the width/2) from individuals located on left of the cline in the sheltered area (in blue) and on the right of the cline in the exposed area (in yellow), and all show a clear bimodality of each parameter. In b. and c., dots are coloured by the shell size gradient, and in d. dots are coloured by the observed heterozygosity (Hobs).

### Bimodal hybrid zones between ecotypes

The high transect-wide LD (Figure S11), the similar cline shapes among inversions (Figure 2a) and the absence of later generation recombinant hybrids (Figure 2b) suggest that the inversions do no behave independently. This led to the question whether the inversions remain coupled in the centre of the hybrid zone. We used cline analyses to compare a unimodal distribution of phenotypes or hybrid indexes in the centre of the hybrid zone to a bimodal or trimodal distribution, putatively maintained by coupling among inversions. We found evidence for bimodality in all three parameters explored: shell size, PC1 score based on the genome-wide polymorphism, hybrid index based on the inversion genotypes (histograms in Figure 2b-d). For each transect, all clines followed similar trends, being centred at 64m and 122m in the northern and southern transect, respectively. In each case, multimodality was maintained in the centres of the clines (red group in histograms Figure 2b-d, and Figures S12 to S17), with bimodal clines often providing similar fits to trimodal clines in the southern transect while trimodality was always the favoured model in the northern transect (Table S3 to S5). The higher number of hybrids found in the northern transect explains why trimodality was more often supported in this transect, but otherwise the two transects were very similar. This multimodality shows that the *L. fabalis* hybrid zones correspond to overlapping lineages maintaining their phenotypic and genetic integrity despite the occurrence of early-generation hybrids with intermediate phenotypes in the centres of the clines. In agreement with this pattern, we inferred that the LD between the differentially fixed inversions was six and 13 times higher in the northern and southern transect (Table S11), respectively, than what was predicted based on the interaction between selection, dispersal and recombination (Detailed method in MatSup p29). Overall, this strong association suggests a strong coupling of the inversions maintaining the bimodal hybrid zone. Interestingly, bimodality was also inferred in the colinear part of the genome when assessed from the PC1 score computed without the inversions (Table S4), or a hybrid index computed with outlier SNPs found outside the inversion (Table S5). Thus, the barrier effect provided by the coupling of the inversions appears to extend into the colinear part of the genome.

### On the origin of the hybrid zone in *L. fabalis*

We used demographic inferences to explore the divergence history of the ecotypes in *L. fabalis*^38^. We asked whether the divergence between ecotypes was better explained by a model of divergence in the face of gene flow (primary divergence), or was the result of a demographic history involving a period of isolation prior to the current contact. We found support for secondary contact in all inferences conducted (using all SNPs, using collinear SNPs and using inversion SNPs only; Figure 3 & Table S8). The inferences made with collinear or inversion SNPs alone supported an ancestral population expansion and a heterogeneous effective population size along the genome (SC_ae_2N). The best models differed mainly by the inferred effective population sizes and effective gene flow, which were an order of magnitude larger in the collinear model than in the inversion model (Table S8 & S9). The relative timing of splits and secondary contacts was similar between collinear and inversion SNP analyses (T_SecondaryContact_/T_Split_ ∼ 0.1, Table S8), but the secondary contact period was shorter in the overall dataset (T_SecondaryContact_/T_Split_ ∼ 0.01). Only the inferences made from the overall dataset showed support for heterogeneous gene flow (best supported model = SC_2N_2M). We inferred 33% of markers experienced a barrier effect where gene flow was reduced by a factor of 10 compared to the remaining background (Table S8).

**Figure 3:**
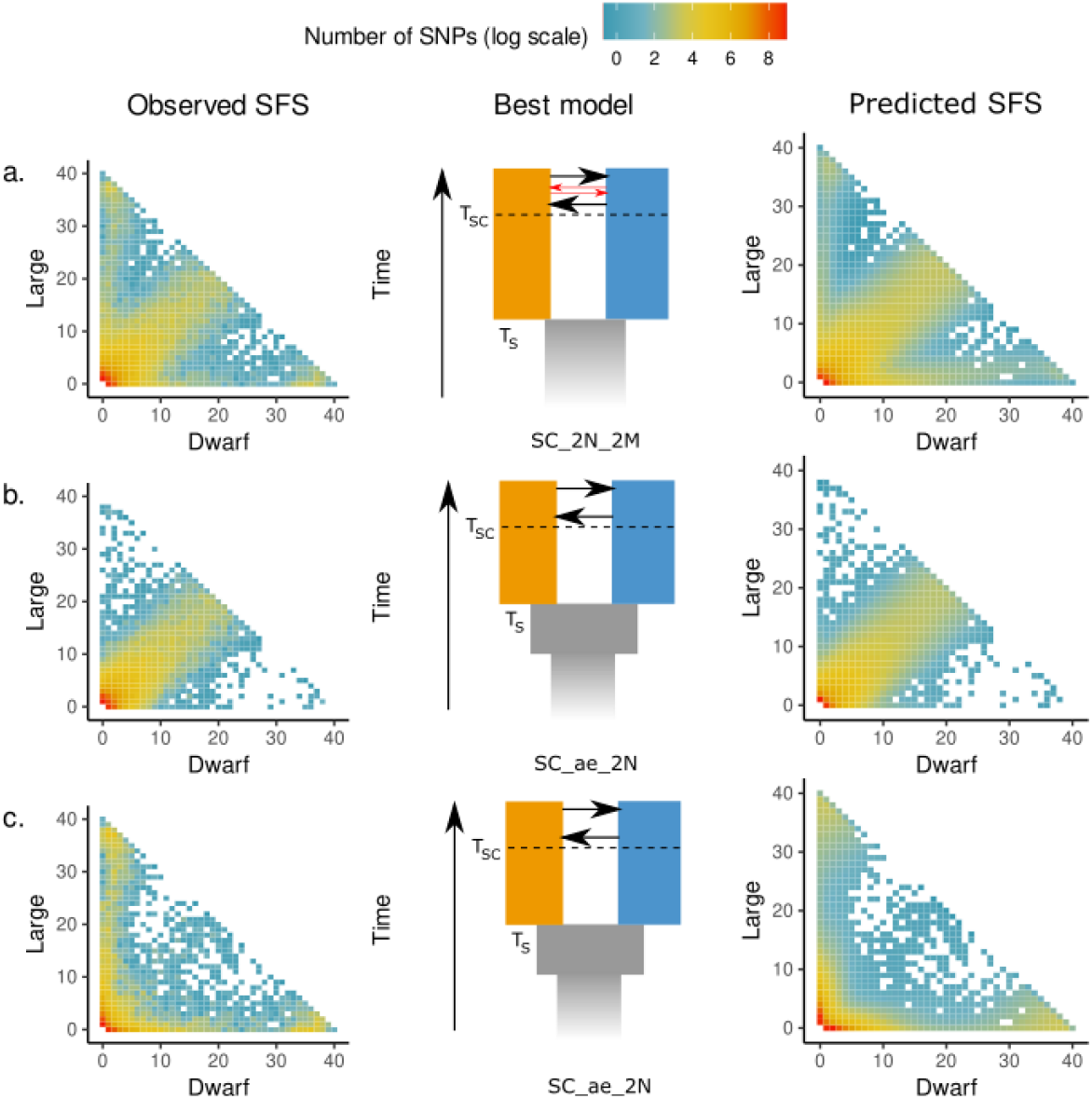
Summary of the demographic inferences performed on the overall dataset (a.), on SNPs located outside the putative inversions (b.) and on SNPs located within the inversions (c.). In each part, the graphs show the observed joint Site Frequency Spectrum (SFS), a schematic representation of the best model inferred based on AIC, and the predicted SFS obtained from the optimisation of the best model. All inferences showed support for a secondary contact (SC) scenario, including heterogeneous effective population size along the genome (2N). Only the overall dataset showed support for heterogeneous gene flow (2M), illustrated by the red arrows in the model depicted in a, while the two other inferences also showed support for population expansion in the ancestral population (ae).

### Correlated landscapes of differentiation across *Littorina* species

The WGS data from *L. fabalis* were analysed using the same reference genome as used in previous studies of *L. saxatilis*^26^, and this allowed us to search for similarities in the architecture of RI. We found a weak but significant correlation in the mean *F*_ST_ per contig between ecotypes across the two species (r²=0.211, p<0.001). This correlation was mostly driven by two linkage groups (LG6 and LG14) carrying an excess of contigs that differed between the contrasting ecotypes in both species (mean *F*_ST_ > 0.1) (Figure 4a). Removing those two LGs from the correlation lead to a correlation coefficient close to 0 (r=-0.004, p= 0.001). The shared differences were clustered at one end of LG6 and LG14 (Figure 4b,c). The genetic signatures in LG14 and LG6 matched those expected from chromosomal inversions (*L. fabalis*: Figures S2 to S10, *L. saxatilis*^21^), suggesting that two chromosomal rearrangements cover similar genomic regions differentiating ecotypes in both species. Indeed, these two putative inversions show clines over ecotype contact zones in both species (*L. fabalis*: Figure 2a, *L. saxatilis*^24,29^). Overall, these results show that the same regions, with similar structural variants, repeatedly contribute to ecotype evolution in distinct, albeit closely-related, species.

**Figure 4:**
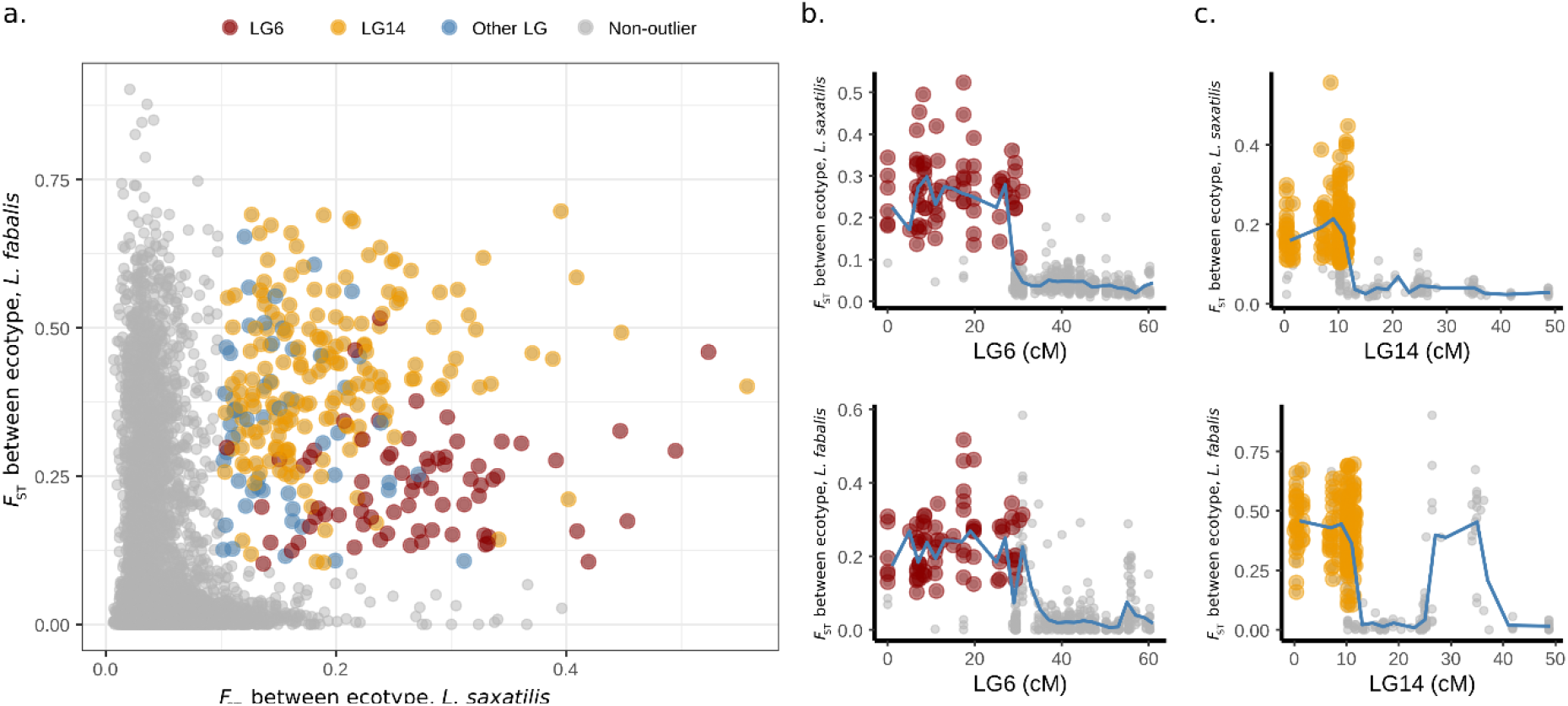
Correlated genomic landscapes of ecotype differentiation in *L. saxatilis* and *L. fabalis*. a. shows the correlation of the mean *F*_ST_ value per contig between ecotypes in both species (r=0.211, p<0.001). The colours highlight contigs with *F*_ST_ > 0.1 in both species. Red dots correspond to shared outliers located on LG6, orange on LG14, and blue on other LGs. c and d. show the variation of mean *F*_ST_ in *L. saxatilis* (top) and *L. fabalis* (bottom) along the two linkage groups carrying a high number of contigs with *F*_ST_ > 0.1 in both species (LG6 and LG14), with dots showing mean *F*_ST_ per contig, and blue line the average *F*_ST_ per cM. The shared outlier contigs from LG6 and LG14 highlighted in a. are highlighted with the same colours in b. and c.

## Discussion

Size is the only major phenotypic difference between the dwarf and the large ecotype of *L. fabalis*^33,39^. In addition, they have slightly different association with wave exposure: the dwarf ecotype is distributed along the more sheltered part of the shore and the large ecotype in the moderately exposed parts, with shifts from one ecotype to the other sometimes taking place over a few meters only^32^. Here we found that the two ecotypes show strong but heterogeneous genomic differences concentrated in 12 putative inversions distributed over nine of the 17 *Littorina* chromosomes. The inversions form steep and coupled clines across hybrid zones between the two ecotypes and, at the contact, a deficiency of heterokaryotypes for the inversions is associated with bimodality of phenotype and genotype distributions. Our demographic analyses strongly suggest that this hybrid zone is a result of secondary contact following a period of isolation. This isolation may have facilitated both adaptive divergence and the accumulation of incompatibilities that decrease hybrid fitness^13^.

The arrangements of nine of the 12 inversions being near fixed different between the two ecotypes results in a strong but highly heterogeneous barrier to gene flow. The strength of this barrier depends on the fitness effects of the barrier loci (i.e., the linked effects of the loci inside each inversion) and the distribution of the loci along the genome^15,40^. We found that strong selection maintains the repeated and highly predictable association between the wave-exposure gradient and the distribution of the two ecotypes, despite the apparently subtle environmental transition. The overall selection involves an extrinsic component: an earlier study using reciprocal transplants and tethering of snails in the seaweed, or on the rock underneath, showed differential survival of the ecotypes with the large ecotype having highest survival in moderately exposed environments, because it better resisted crab attacks if dislodged from the seaweed during heavy wave action^41^. Bimodality in the contact zone might also imply assortative mating^9^, although previous field observations suggested close to random mating^32^, arguing against this possibility. Yet, we found evidence that the association between arrangements of different inversions was much stronger than expected from the effect of selection acting independently on each inversion. These results, together with the deficiency in heterokaryotypes, imply coupling between inversions, perhaps due to epistatic effects with extrinsic and/or intrinsic components. This coupling of multiple inversions may result in an accumulation of underdominant effects that contributes to the absence of recombinant hybrids and so strengthens the overall barrier to gene flow between *L. fabalis* ecotypes. It also implies that *L. fabalis* ecotypes are further along the speciation continuum than are the Swedish *L. saxatilis* ecotypes.

The structure of the hybrid zones in *L. fabalis* differs from the situation in *L. saxatilis* where the coupling of the different inversions is weak in the centres of the hybrid zones^42^, and almost all inversions remain polymorphic at cline ends^21,24,29^. These differences coincide with different evolutionary histories of the ecotypes in the two species. While the unimodal hybrid zones between the crab and wave ecotypes of *L. saxatilis* have evolved by divergent selection with short interruptions to gene flow, at most ^43^, we found that the *L. fabalis* hybrid zones originated from a secondary contact after a long period of isolation. The inferred period of separation was during the Quaternary glacial period (68-150k year ago) while the secondary contact matches the post-glacial period (2-9k year ago). The allopatric divergence of *L. fabalis* ecotypes likely facilitated the coupling of barriers to gene flow, including components of adaptation and genetic incompatibility, which is now involved in the maintenance of the bimodal hybrid zones.

Despite the differences in the hybrid zone structures, and the different demographic backgrounds, the evolution of *L. fabalis* and *L. saxatilis* ecotypes shares some interesting features. For instance, the barriers to gene flow focus on many polymorphic inversions, some of which remaining highly polymorphic within ecotype in both species. Such a spatial pattern is frequently found in the inversions of *L. saxatilis*^24,29^, and in other systems^19,22^. These polymorphisms could be driven by some type of balancing selection maintaining the two arrangements within one ecotype^44^, such as frequency dependence or heterokaryotype advantage. Furthermore, our comparative analysis suggests that two inversions, on LG6 and LG14, cover similar genomic regions in the two species. In Swedish populations of *L. saxatilis*, these are the two most important inversions involved in ecotype differentiation^26,29^ and they host many QTL associated with phenotypic traits under divergent selection in this area^27,28^. These inversions could have the same origin and be shared by a recent introgression event between *L. saxatilis* and *L. fabalis*, or through repeated selection of an ancestral polymorphism segregating in the standing variation of both species. Alternatively, the repeated and independent origins of new inversions in the same genomic regions in both species, presumably favoured by selection because they captured co-adapted alleles, could also have led to this correlated genomic landscape of ecotypic differentiation. Correlated landscapes of differentiation between sister taxa are present in other systems^45–47^, and can often be explained by the effect of background selection on shared recombination landscapes^48^. The chromosomal inversions detected here represent a low recombining region expected to amplify the hitchhiking effects associated with background selection. However, in the snails, these regions are associated with sharp allelic clines in both species (here and in^24,29^), which suggests that shared architectures of barriers to gene flow can also lead to correlated landscapes of differentiation.

In *L. fabalis*, the barriers provided by the coupling of inversions extend into the colinear part of the genome, but the reduction in gene flow in these collinear parts, maintained through multiple generations of backcrosses, only marginally affects the background differentiation. Indeed, the collinear *F*_ST_ value of 0.03 between the *L. fabalis* ecotypes is within the range of *F*_ST_ values found between the collinear regions of separate *L. saxatilis* ecotypes with distribution patterns across similar types of wave exposure gradients^29^. This contrast between rearranged and colinear differentiation is similar to what has been found in other systems, such as in migratory vs stationary ecotypes of cod^20^, meadow vs forest ecotypes of deer mice^19^, and dune vs field ecotypes of sunflower^22^. Indeed, restriction of recombination in inversion heterozygotes increases recombination in the collinear genome^49^, accentuating the difference in barrier strength experienced by neutral alleles, and so the observed differentiation between inverted and collinear parts of the genome. Such increased recombination in the collinear regions might lead to stronger migration load^50^, and impede the formation of genome-wide barriers to gene flow. Overall, while our results concur with previous work highlighting the importance of large chromosomal rearrangements for ecotypic divergence and barriers to gene flow, and show that the coupling of several inversions can generate strong reproductive barrier, it remains an open question whether this type of genomic architecture of reproductive isolation will eventually lead to the completion of the speciation process.

## Material and methods

### Geographic and genomic sampling

Snails were collected over two transects covering a wave exposure gradient located along the southern and northern shores of Lökholmen island on the Swedish west coast (58°88N 11°11E). The position and elevation of each snail along the transects were recorded with a Trimble total station. The three-dimensional position of each snail was transformed into one dimensional spatial coordinates using least-cost distance following Westram et al^29^. We took pictures of each snail with the aperture up alongside a scale, and the size of each snail was measured in ImageJ as the longest distance from the aperture rim to the apex. Snails were thereafter dissected and a piece of foot tissue was stored in 95% ethanol. We extracted DNA from tissue samples using the protocol by Panova et al^51^. After measuring concentrations and purity of the extractions on Qbit and nanodrop, DNA was shipped to SciLifeLab (Sweden) for individual Nextera WGS library preparations, followed by sequencing on a NovaSeq S6000. Targeted coverage was 5X except for 12 individuals taken from the ends of the transects which were sequenced at ∼15X coverage.

### Bioinformatics pipeline

The WGS short reads were mapped to the reference genome of the related species *L. saxatilis* using *bwa-mem*^52^. This reference genome is fragmented but is complemented by a dense linkage map^24^. Mapped reads were processed though the GATK pipeline^53^ following good practice guidelines including the removal of PCR duplicates and base-pair recalibration. Further filtration steps were applied using vcftools^54^ in order to keep only bi-allelic SNPs sequenced in at least 90% of the individuals and with an average minimum sequencing depth of 3 and maximum of 15 across samples. This raw dataset contained 1.38M SNPs, which were further filtered depending on the analyses.

### Population structure analyses

For these analyses, the raw dataset was thinned by keeping only SNPs with a minor allele frequency above 5%, leading to 451k SNPs. This data set was further pruned for physical linkage by keeping 1 SNP per 1k base-pair chosen at random (based on LD decay in *Littorina*^24^), resulting in a total of 110k SNPs. We computed a PCA on all individuals (both transects included) from the 110k SNPs pruned for linkage using the R package adegenet^55^. Pairwise *F*_ST_ values between ecotypes were calculated per pruned-SNP using the 20 snails nearest the ends of both transects, analysed independently for each transect using vcftools^54^, and visualized using Manhattan plot in ggplot. Then, we used a sliding window approach over 1cM bins using the 458k SNPs unpruned for linkage to compute local PCA with adegenet^55^ and variation of heterozygosity with the *Hsplit* function (Reeves et al. in prep) to explore the variation in structure and diversity patterns along the *L. saxatilis* linkage map^24^. Pairwise LD between SNPs was then calculated per linkage group using only markers with minor allele frequency above 25% using LDheatmap^56^. Based on variation patterns of LD, *Hsplit, F*_ST_ and local PCA along the chromosomes, we visually defined 14 linkage blocks carrying high differentiation between ecotypes, which were likely due to chromosomal rearrangements. Then, we used unlinked SNPs from outside the linkage blocks to compute the relatedness among all pairs of snails using the *a*_*jk*_ statistic from Yang et al.^57^ in vcftools^54^. These analyses allowed us to identify multiple pairs of full-sibs and half sibs, which we used to calculate the standard deviation of the dispersal distance between parent and offspring (σ) from the mean transect distance between sibling individuals 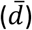 using the following formula:*σ* = 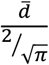 More details about this equation and its assumptions are available in supplementary p28. We computed DAPC using SNPs from within each of the linkage blocks for a k value of three clusters using adegenet^58^ in order to infer the karyotypes of the putative chromosomal rearrangements for each snail. These karyotypes were finally used to calculate the hybrid index and the observed heterozygosity

### Cline analyses

Cline analyses were performed within a Bayesian framework using the R package Rstan^60^. We fitted cline models for continuous traits to the variation observed along the two transects in shell length, the PC1 score computed for the overall dataset, the PC1 score for the dataset from SNPs outside chromosomal rearrangements, and the hybrid index computed from the chromosomal rearrangement genotypes. We compared the ability of 5 contrasted cline models to fit the data, following the formulation in the Cfit package^8^ but re-coded in custom Rstan functions: a unimodal cline, two bimodal clines, one with and one without introgression, and two trimodal clines, one with and one without introgression. Unimodal clines are generally used for quantitative traits when the phenotype observed in the centre of the hybrid zone follows a normal distribution centred around the mean of the two parental phenotypes while multimodal clines are expected when the parental phenotypes remain more or less discrete even in the centre of the hybrid zone. Bimodal clines are expected between species overlapping in the centre of a hybrid zone that maintain phenotypic differences in the centre of the cline similar to those between the extremes. Such hybrid zones are expected at strong RI or from traits with a simple genetic basis or threshold behaviour. Furthermore, the phenotype can also be inferred as trimodal if hybrids with intermediate phenotypes occur in the centre of the cline but have low fitness, avoiding production of a hybrid swarm. The differences between the parental modes near to the centre of the cline can be reduced in the case of introgression, which is the reason why the introgression was also modelled in the multimodal clines. Chromosomal rearrangement frequency clines were also fitted in Rstan using the karyotypes inferred from the local DAPC. The cline functions tested included the estimation of F_IS_, assumed to peak at the cline centre and estimated at 5 points, following the formulation in Cfit^8^ but also rewritten as Rstan functions. Finally, the coefficients of selection required to maintained the cline were calculated from the width of the cline, and the estimate of dispersal, σ, inferred from the distance between pairs of related individuals, using the formula from Barton and Gale^6^ assuming an abrupt habitat change and no dominance effect: 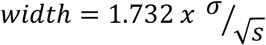 where *s* is the reduction in mean fitness at the cline centre.

Furthermore, we explored the allelic clines at each SNPs from the linked dataset (i.e. not filtered to one SNP per kb) with a MAF above 0.3 and a difference in allele frequency between the extreme parts of a transect above 0.1. In total, we inferred simple sigmoid clines, without assuming fixation at the ends, from 90k SNPs using maximum likelihood in bbmle (which is less computationally intensive than Rstan). Then, following the procedure from Westram et al.^24,29^, we compared the AIC of model with fixed frequency and linear frequency variation to clinal variation along the transect. Clinal fits showing an AIC difference >4, compared with a non-clinal fit, were considered as significant clines. The goodness of each cline fit was then estimated using a generalized linear regression model with binomial error to evaluate how well frequencies estimated from the cline explained the genotype observed for each snail given their position on the transect.

### Demographic analyses

We used the software *moments*^61^ to infer the demographic history of divergence between the large and the dwarf ecotype of *L. fabalis*. This software uses the information contained in the joint Site Frequency Spectrum to fit contrasted demographic scenarios using an approach based on a diffusion approximation. We calculated the SFS using 20 snails sampled from each end of each transect. To avoid bias in the SFS due to the miss-calling of the heterozygotes with low coverage depth in our data, one allele per polymorphic position was randomly sampled independently in each individual. Individuals from both transects were then pooled to produce a SFS of 40 chromosomes from the large ecotype against 40 chromosomes (i.e. one chromosome per individual) from the dwarf ecotype and to explore the divergence between ecotypes (“overall SFS”). This SFS was further subsampled to include only the SNPs from within the putative chromosomal inversions (“inversion SFS”) or from outside the inversions (“collinear SFS”). Demographic inferences were then made independently for the three SFS. In total, we compared the ability of 28 demographic scenarios to reproduce the observed SFS^35,62^, including scenarios of allopatric divergence (4 variants of a strict isolation (SI) model), and divergence with gene flow (8 variants of an isolation-with-migration (IM) model, 8 variants of an ancestral migration (AM) model, and 8 variants of a secondary contact (SC) model). All scenarios were adjusted to the observed data using the optimisation procedure described in Portik et al^63^. The different demographic scenarios were compared using AIC, with models differing in AIC by <10 considered as equally good. Demographic parameters were transformed using the method described in Rougeux et al^64^, using a mutation rate of 1 x 10^−8^ per bp and generation and a generation time of 1 year^65^.

### Interspecific comparative analysis of differentiation

As the same reference genome was used between this work and previous work performed on ecotype evolution in *L. saxatilis*, we could compare the genomic landscape of ecotype differentiation between the two species. We used a simple correlation test on mean *F*_ST_ per contig between ecotypes in each species using SNPs derived from the whole genome, and using SNPs from each linkage group independently. Those *F*_ST_ values were calculated independently in both species, using the lcWGS data described in the present study for *L. fabalis*, and previously published *F*_ST_ values from Morales et al^26^ computed from pool-seq WGS data for *L. saxatilis*. For L. saxatilis, we used only the *F*_ST_ between ecotypes from two sites (ANG and CZB), which were located on two islands of the same archipelago as the *L. fabalis* hybrid zone studied here. Shared outlier regions were then considered as any contig with mean *F*_ST_ between ecotypes above 0.1 (arbitrary threshold) and above the 5% quantile in both species, as well as above the 1% quantile in each species.

