## Supplementary Material for "Coupling of 12 chromosomal inversions maintains a strong barrier to gene flow between ecotypes"

Table S7: Summary table of the demographic inferences ...................................... 1

### Signature of chromosomal rearrangements

#### Table S1: Summary statistics of the putative inversions.

In order of appearance, the table shows the name of the inversion, the size in base pairs, the size in the linkage map (in cM), the total number of SNPs, the number of SNPs with minor allele frequency (maf) above 5%, the average $F_{ST}$ between homokaryotypes, the average LD among SNPs with maf >0.3 and with 1 SNPs per kb (all individuals from both transects were pooled), the number of SNPs that are fixed different between homokaryotypes, the proportion of differentially fixed SNPs among those with maf >5%, the number of SNPs with $F_{ST}$ > 0.9 between homokaryotypes, and the proportion of SNPs with maf > 5% and $F_{ST}$ > 0.9.

| Inversion | Size (Mbp) | Size (cM) | $n^{SNPs}$ | $n^{SNPs>5\%}$ | $F_{ST}$ | LD | $n^{fixed}$ | $P^{fixed}$ | $n^{FST>0.9}$ | $P^{FST>0.9}$ |
|---|---|---|---|---|---|---|---|---|---|---|
| LG3_inv1 | 33,160 | 47,034 | 110996 | 43242 | 0,43 | 0,425 | 650 | 0,015 | 7078 | 0,164 |
| LG4_inv1 | 8,000 | 1,831 | 21177 | 8929 | 0,452 | 0,461 | 255 | 0,029 | 1972 | 0,221 |
| LG4_inv2* | 8,330 | 14,666 | 29666 | 12002 | 0,43 | 0,433 | 384 | 0,032 | 2562 | 0,213 |
| LG4_inv3* | 5,171 | 15,004 | 8393 | 3324 | 0,328 | 0,262 | 39 | 0,012 | 465 | 0,14 |
| LG6_inv1 | 10,510 | 33,837 | 30691 | 11316 | 0,331 | 0,244 | 180 | 0,016 | 1319 | 0,117 |
| LG6_inv2 | 2,206 | 0,624 | 7146 | 2287 | 0,205 | 0,106 | 29 | 0,013 | 181 | 0,079 |
| LG7_inv1 | 4,285 | 5,546 | 12758 | 4445 | 0,306 | 0,284 | 74 | 0,017 | 553 | 0,124 |
| LG7_inv2 | 5,568 | 12,687 | 21297 | 7901 | 0,304 | 0,273 | 112 | 0,014 | 900 | 0,114 |
| LG8_inv1 | 2,715 | 5,328 | 5660 | 2024 | 0,236 | 0,267 | 7 | 0,003 | 102 | 0,05 |
| LG11_inv1 | 4,409 | 38,062 | 11930 | 4926 | 0,412 | 0,345 | 119 | 0,024 | 756 | 0,153 |
| LG13_inv1 | 10,147 | 35,951 | 26761 | 9956 | 0,327 | 0,279 | 67 | 0,007 | 848 | 0,085 |
| LG14_inv1[+] | 12,293 | 11,700 | 37631 | 14918 | 0,472 | 0,489 | 360 | 0,024 | 3168 | 0,212 |
| LG14_inv2[+] | 1,336 | 9,893 | 2808 | 1043 | 0,524 | 0,508 | 66 | 0,063 | 333 | 0,319 |
| LG16_inv1 | 5,192 | 31,229 | 9869 | 3351 | 0,299 | 0,294 | 14 | 0,004 | 219 | 0,065 |

#### Table S2: Summary statistics of the three karyotypes of each putative inversion.

The karyotypes of each inversion were inferred from the DAPC analyses presented in Figure S5. The most frequent arrangements among dwarf ecotypes are labelled "D", the most frequent arrangements among large ecotypes are labeled "L" and heterokaryotypes are labelled "D/L". For each karyotype of each inversion, the table summarizes the number of individuals, the average shell size of the snails (Size), the mean observed heterozygosity (Hobs); the proportion of SNPs with Hobs > 0.5 ($P_{SNPs}$ Hobs >0.5), and the mean LD between SNPs within each karyotype.

| Inversion | N | | | Size | | | Mean Hobs | | | $P_{SNPs}$ Hobs >0.5 | | | LD within karyotype | | |
|---|---|---|---|---|---|---|---|---|---|---|---|---|---|---|---|
| | D/D | D/L | L/L | D/D | D/L | L/L | D/D | D/L | L/L | D/D | D/L | L/L | D/D | D/L | L/L |
| LG3_inv1 | 123 | 54 | 118 | 8,59 | 10,52 | 11,49 | 0,17 | 0,37 | **0,12** | 0,02 | 0,33 | 0,02 | 0,01 | 0,14 | 0,01 |
| LG4_inv1 | 93 | 61 | 141 | 8,57 | 9,88 | 11,21 | **0,13** | 0,39 | 0,14 | 0,02 | 0,35 | 0,02 | 0,02 | 0,15 | 0,01 |
| LG4_inv2* | 99 | 52 | 144 | 8,53 | 9,83 | 11,28 | **0,14** | 0,38 | 0,15 | 0,02 | 0,32 | 0,02 | 0,01 | 0,14 | 0,01 |
| LG4_inv3* | 99 | 52 | 144 | 8,53 | 9,83 | 11,28 | **0,16** | 0,34 | 0,18 | 0,02 | 0,24 | 0,02 | 0,02 | 0,10 | 0,02 |
| LG6_inv1 | 67 | 93 | 135 | 8,64 | 9,54 | 11,21 | **0,16** | 0,32 | 0,17 | 0,03 | 0,23 | 0,01 | 0,02 | 0,08 | 0,01 |
| LG6_inv2 | 60 | 100 | 135 | 8,63 | 9,82 | 10,97 | 0,18 | 0,27 | 0,19 | 0,02 | 0,13 | 0,02 | 0,02 | 0,07 | 0,02 |
| LG7_inv1 | 107 | 68 | 120 | 8,58 | 10,25 | 11,37 | **0,14** | 0,30 | 0,17 | 0,02 | 0,20 | 0,03 | 0,01 | 0,09 | 0,01 |
| LG7_inv2 | 106 | 55 | 134 | 8,49 | 10,13 | 11,37 | 0,16 | 0,32 | 0,17 | 0,02 | 0,21 | 0,02 | 0,02 | 0,09 | 0,01 |
| LG8_inv1 | 117 | 43 | 135 | 8,60 | 10,25 | 11,35 | **0,16** | 0,27 | 0,17 | 0,03 | 0,14 | 0,03 | 0,01 | 0,11 | 0,01 |
| LG11_inv1 | 61 | 82 | 152 | 8,63 | 9,18 | 11,19 | **0,11** | 0,34 | 0,16 | 0,02 | 0,26 | 0,02 | 0,02 | 0,13 | 0,01 |
| LG13_inv1 | 95 | 64 | 136 | 8,55 | 9,82 | 11,32 | **0,16** | 0,34 | 0,18 | 0,03 | 0,25 | 0,03 | 0,01 | 0,10 | 0,01 |
| LG14_inv1[+] | 115 | 41 | 139 | 8,58 | 9,75 | 11,47 | 0,15 | 0,40 | **0,12** | 0,02 | 0,37 | 0,02 | 0,01 | 0,16 | 0,01 |
| LG14_inv2[+] | 115 | 41 | 139 | 8,58 | 9,75 | 11,47 | 0,15 | 0,44 | **0,13** | 0,03 | 0,42 | 0,02 | 0,01 | 0,20 | 0,02 |
| LG16_inv1 | 113 | 54 | 128 | 8,55 | 10,26 | 11,41 | **0,16** | 0,32 | 0,18 | 0,02 | 0,23 | 0,03 | 0,02 | 0,12 | 0,02 |

#### PCA analyses

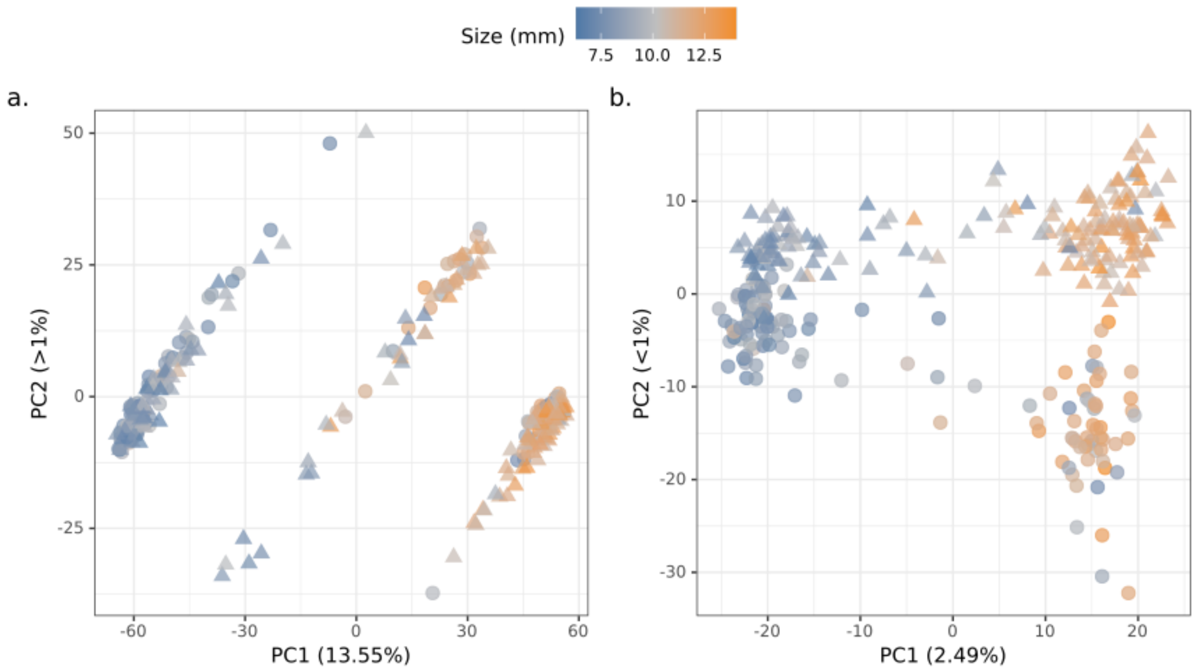

Figure S1: PCA analyses. Each dot is an individual snail coloured by its size according to the color gradient above. Triangles and circles distinguish snails from the northern and southern transects, respectively. In a., the PCA was computed on the overall dataset, and b. without SNPs from inside the LD blocks. PC1, showing 13.55% and 2.49% of the total variation in a. and b. respectively, distinguished snails by size / ecotypes, and PC2 distinguished snails by transect / geography only in b. This geographical structure was more pronounced in large individuals.

#### Linkage disequilibrium

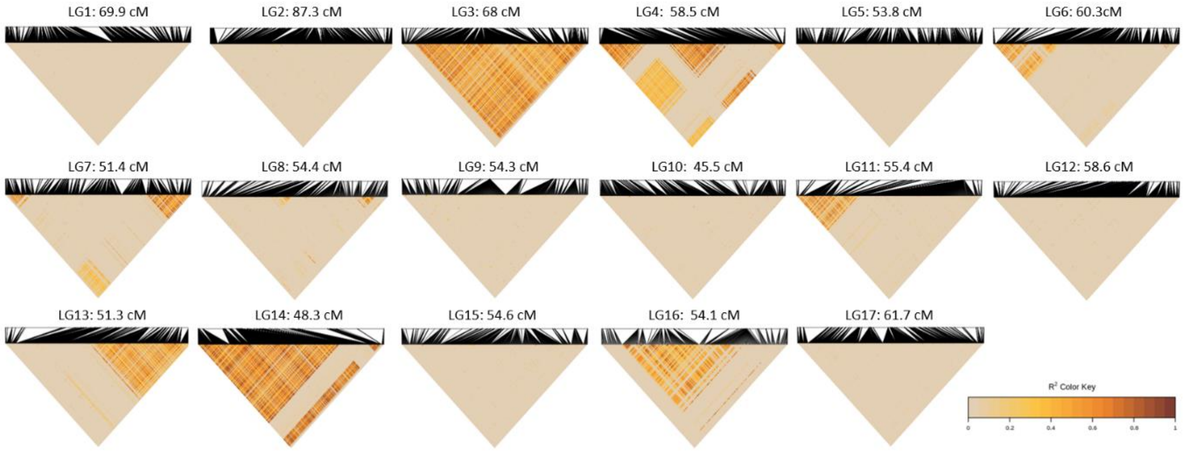

Figure S2: LD heatmap per LG. LD was computed using r² from SNPs with MAF of 30% and minimum distance of 1kb between SNPs using the R package LDheatmap. Pairwise LD value are proportional to the color gradient on the bottom right. Distance between SNPs on the linkage map of *L. saxatilis* (Westram et al. 2018) is represented by the scale on the top of each graph.

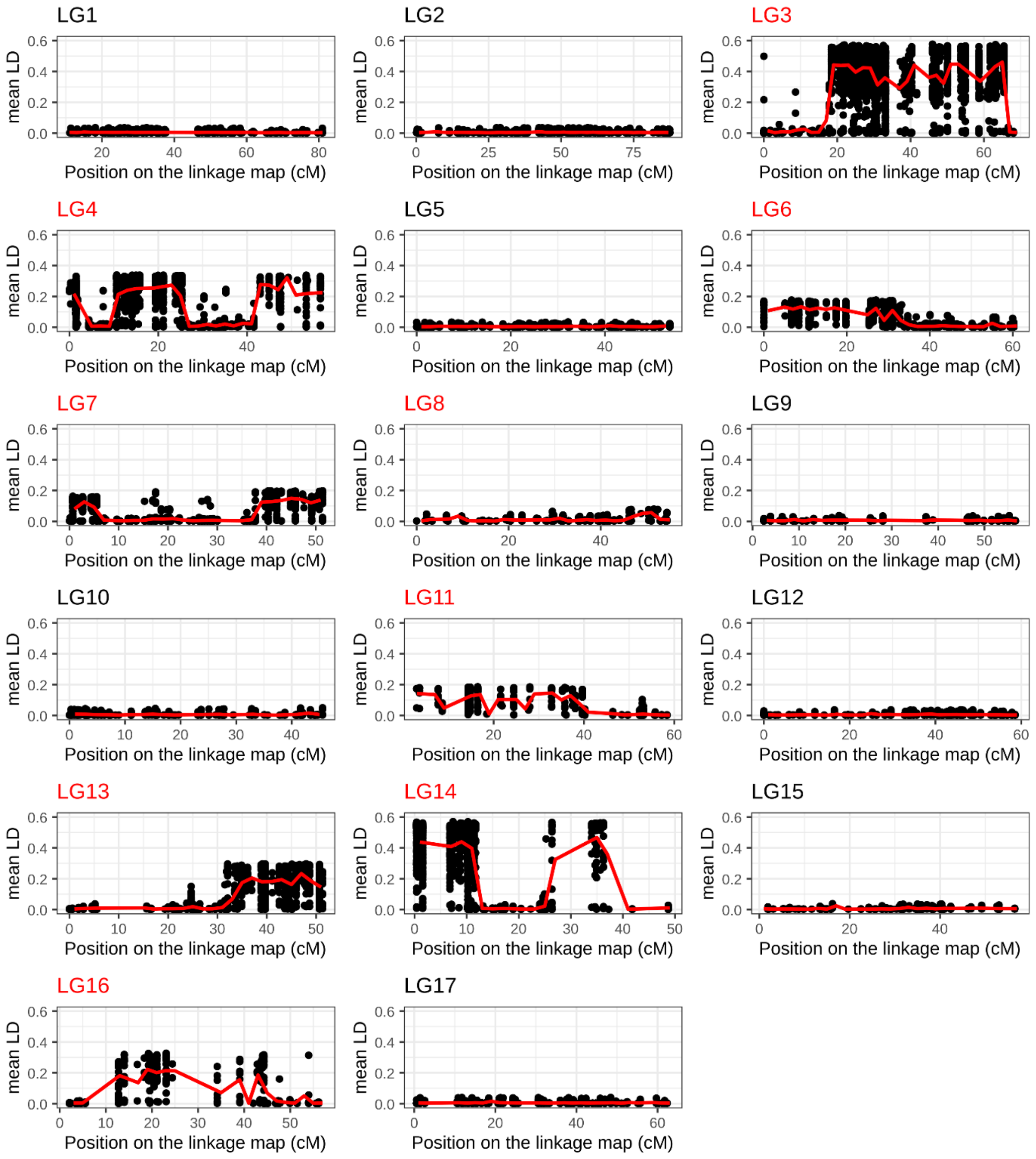

Figure S3: Variation of mean LD along each LG. LD was computed with r² using SNPs with minimum MAF of 30% and minimum distance of 1kb between SNPs using the R package LDheatmap. Each dot represents the mean LD of a given SNP with all other SNPs from the LG and the solid line represent the variation of mean LD along the LG across bins of 2 cM. In this graph, the larger the arrangement is, the higher will be its effect on the mean LD variation along the LG.

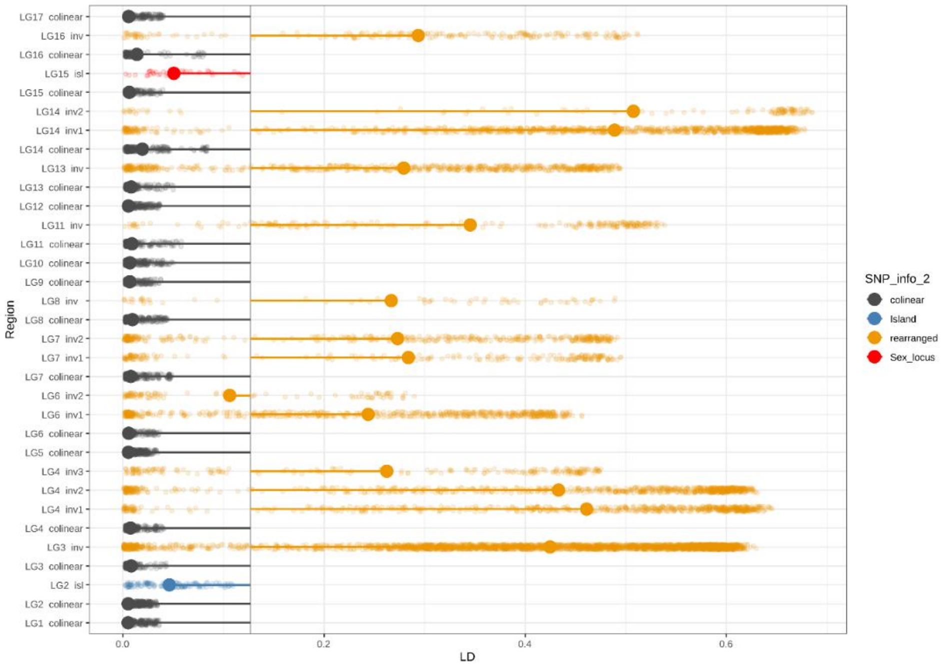

Figure S4: Mean LD per LG over collinear and putatively inverted regions. LD was computed with the r² method with the R package LDheatmap using only SNPs with minimum MAF of 30% and minimum distance of 1kb between SNPs. Average genome-wide LD is represented by the vertical line. The average LD within an inversion (orange) or in collinear region (grey) of a given LG is represented by the large dot, and the deviation from the mean LD is represented by the horizontal line. LD within most inversions (all expect for LG6_inv2) is higher than the average genome-wide value while LD within the collinear regions is lower than the average genome wide. LD within the genomic island of differentiation in LG2 (Figure 1) is represented in blue, and LD within the sex determining region is represented in red.

#### Local PCA and Clustering:

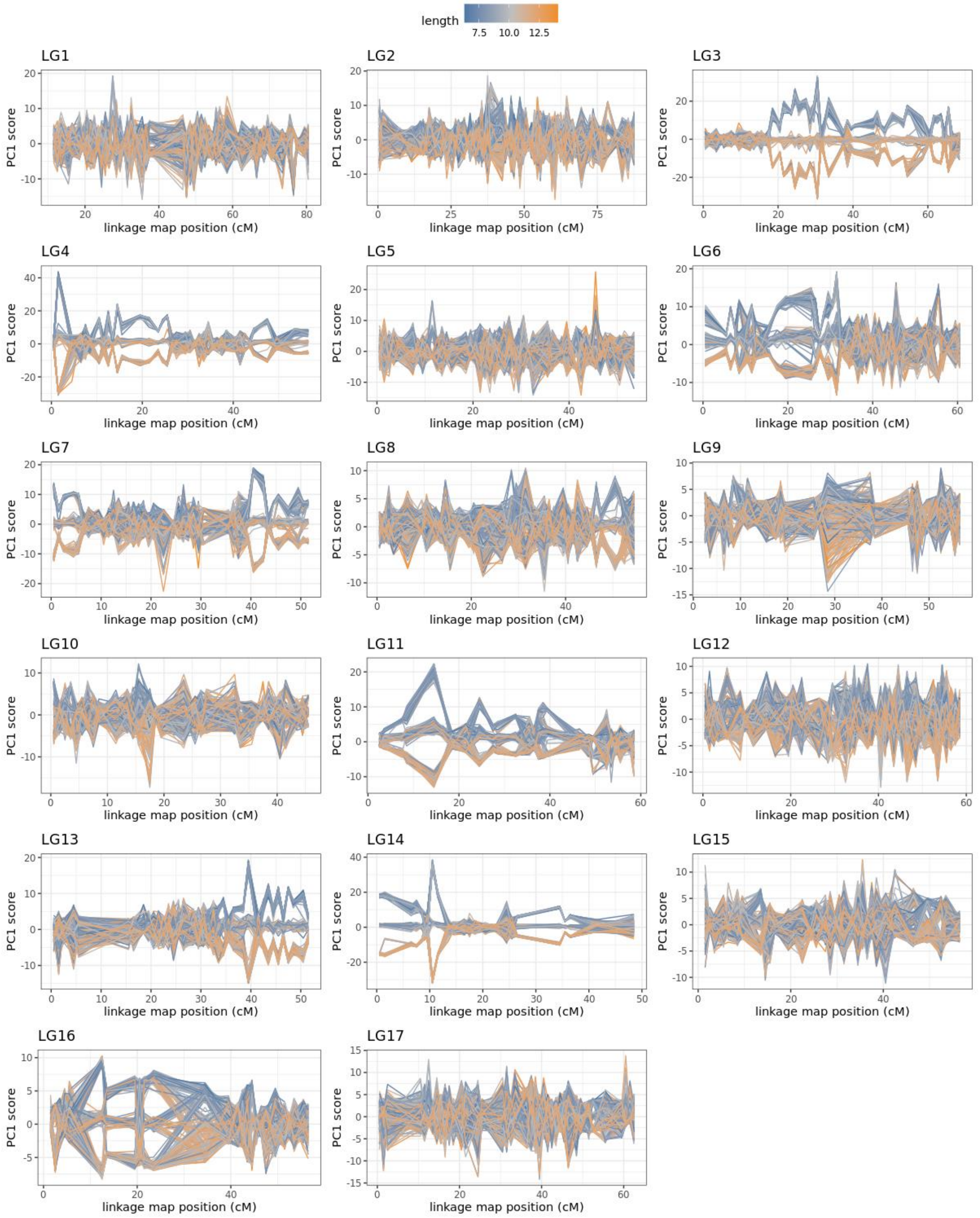

Figure S5: local PCA per bin of one cM across the 17 linkage groups of the *L. saxatilis* linkage map. For each bin of 1cM, the position of the snail along the first axis of the PC was extracted, and plotted against the bin location on the map. Lines were then drawn to link the different PC positions of a single snail across the maps. Thus, each line corresponds to one snail, which was then colored by its size (orange for large, blue for dwarf and grey for intermediate size). The split of the samples into three distinct groups over many consecutive windows of the LG3, LG4, LG6, LG7, LG8, LG11, LG13, LG14 and LG16 is one of the expected signatures from large chromosomal rearrangements, such as inversions.

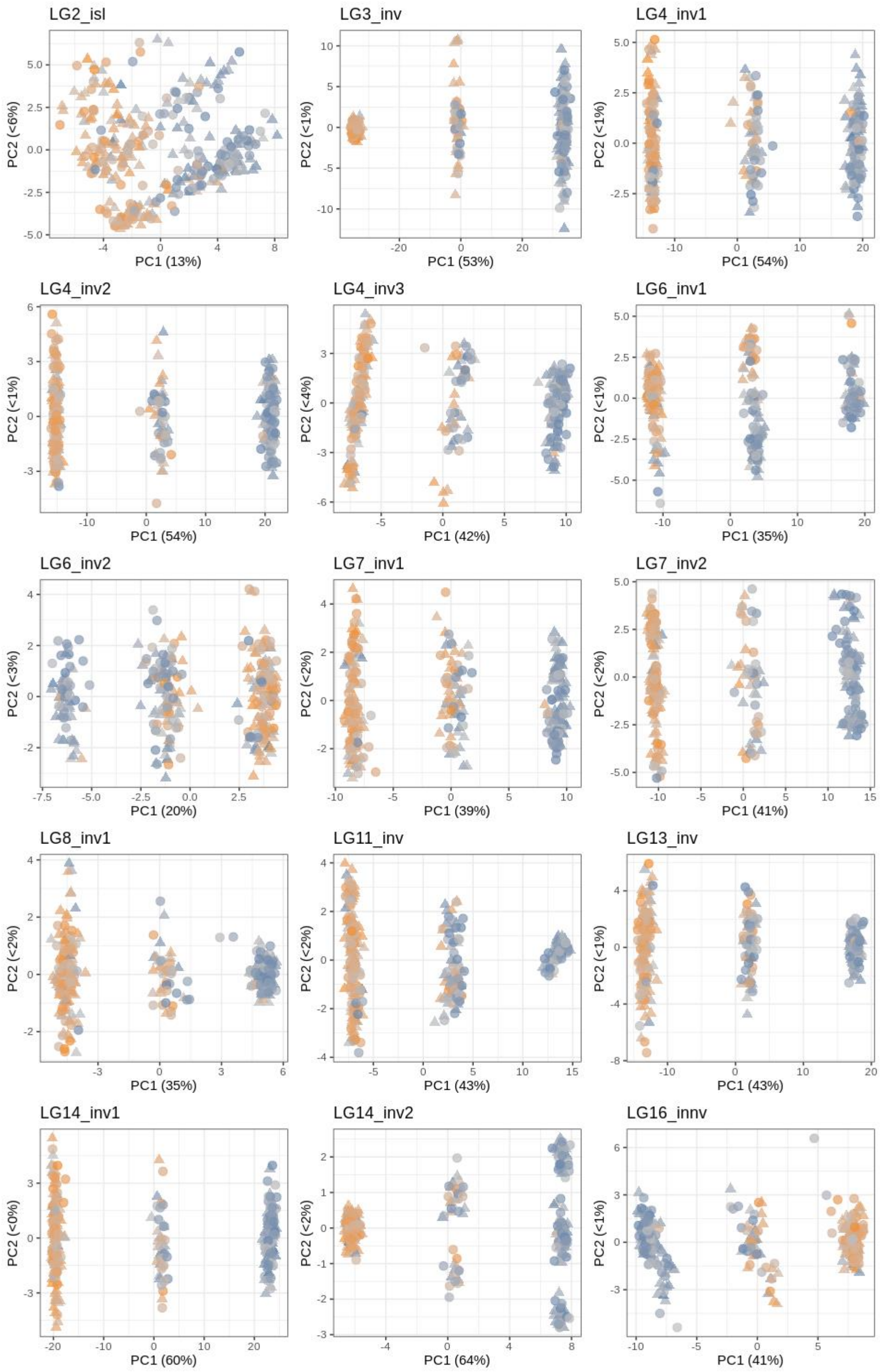

Figure S6: PCA within the main LD blocks. Samples are colored by their size (orange for large, blue for dwarf and grey for intermediate size). Each PCA except the first one ("LG2_isl") shows a clear division of the samples into three groups, with one group located in the center of the two others. This signature is expected from large chromosomal rearrangements, such as inversions.

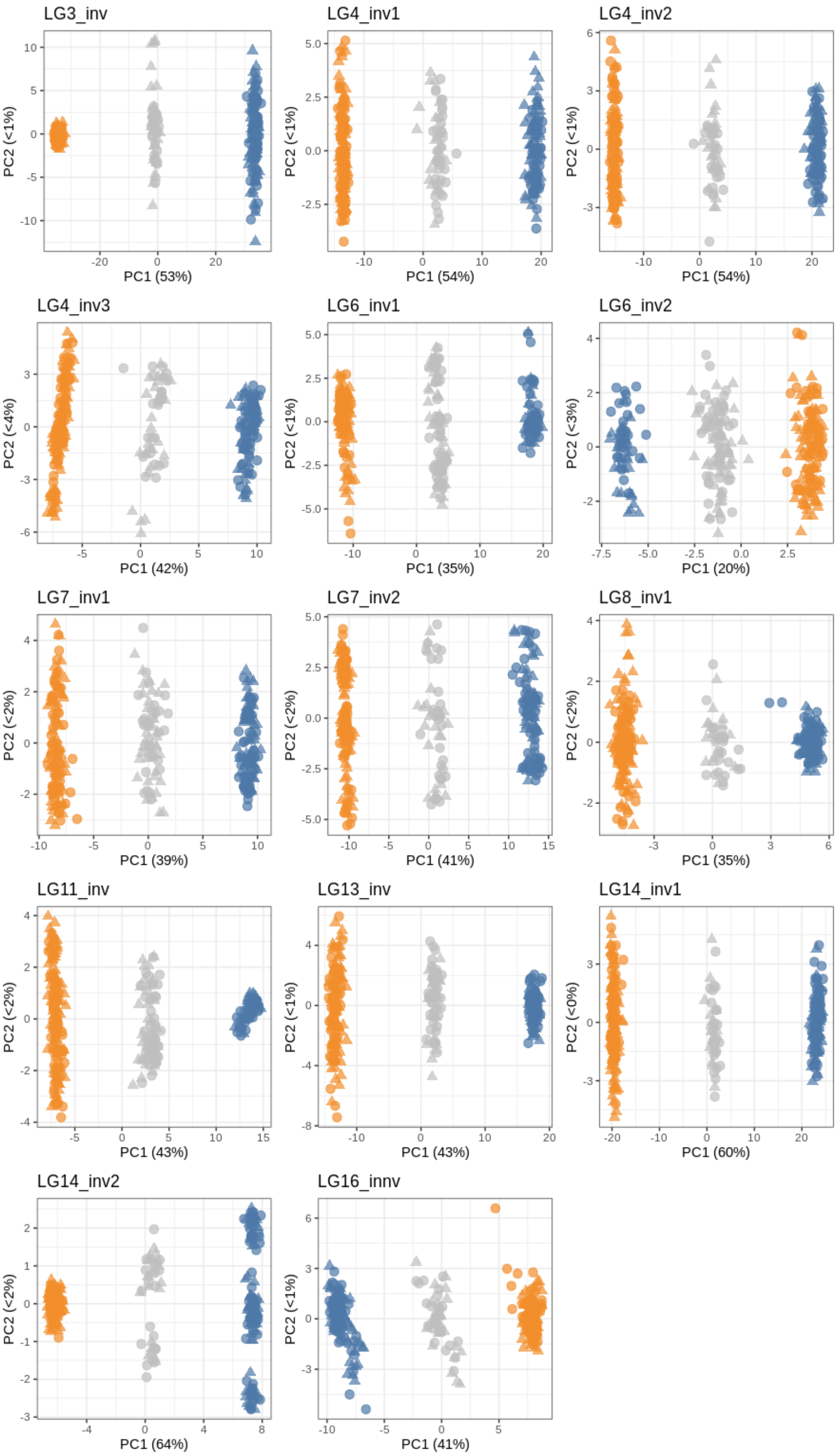

Figure S7: DAPC within the main LD block. The samples are colored by outcome of a clustering analysis performed for k=3 in adegenet. Results of this clustering were used as karyotype input in later analyses, with grey assigned as heterokaryotype (encoded D/L), blue as homokaryotype of the most frequent arrangement in the dwarf ecotype (D/D), and orange as homokaryotype for the most frequent arrangement in large (L/L).

#### Heterozygosity

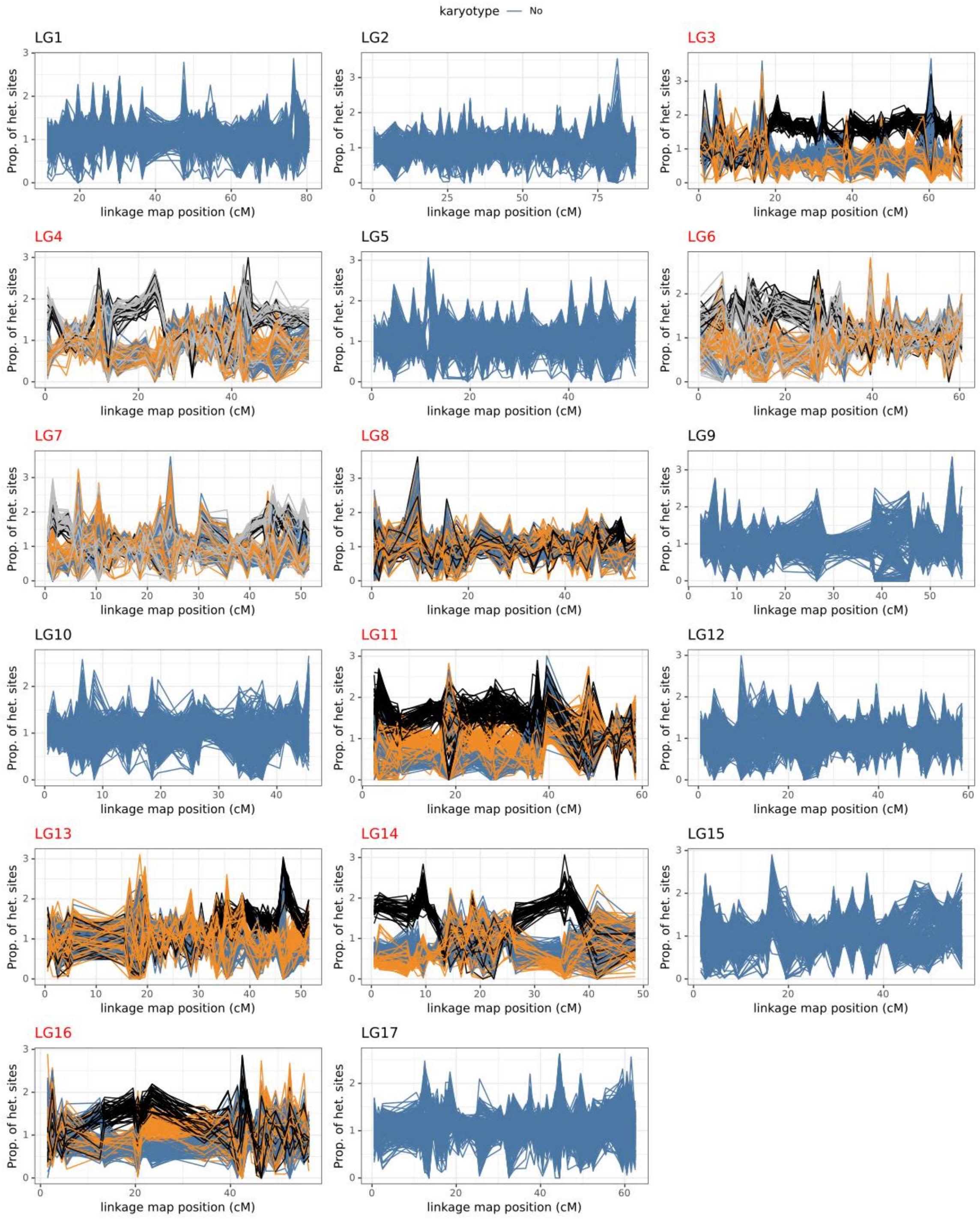

Figure S8: Observed heterozygosity per bin of one cM across the 17 linkage groups of the *L. saxatilis* linkage map. For each bin of 1cM, the proportion of heterozygous sites was calculated by dividing the number of heterozygous positions by the total number of SNPs in the bin. As this proportion is also affected by the coverage that is quite variable across individuals in our lcWGS, these proportions were standardized by the individual average heterozygosity outside the putative inversions. Thus, for a particular snail, an individual value of 2 in a given window means that there is twice as many heterozygous sites per SNP in the window as in the rest of the genome. Lines were then drawn to show the variation of the standardized proportion of heterozygous sites of a single snail across the LG maps. These lines were colored in blue by default, except for linkage groups with inversions that were colored in blue when the individuals were homozygous D/D, orange for homozygous L/L, black for heterozygous D/L, and grey for chromosomes with two arrangements carrying different karyotypes (in LG4, LG6 and LG7). D/L individuals always show higher heterozygosity over the putatively rearranged genomic regions.

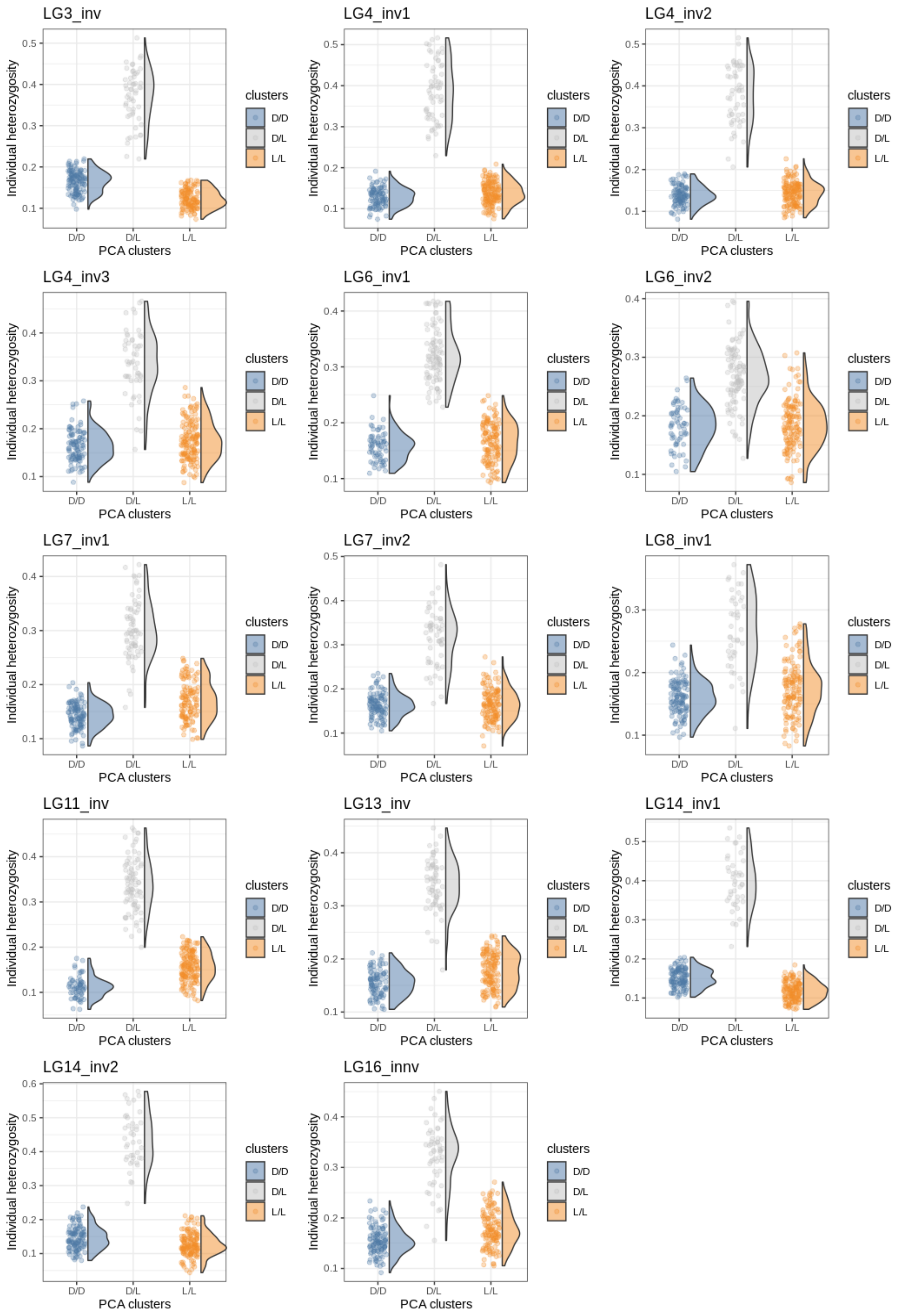

Figure S9: Proportion of heterozygote sites per individual spit according to the DAPC cluster. The cluster D/L always shows the highest level of heterozygosity.

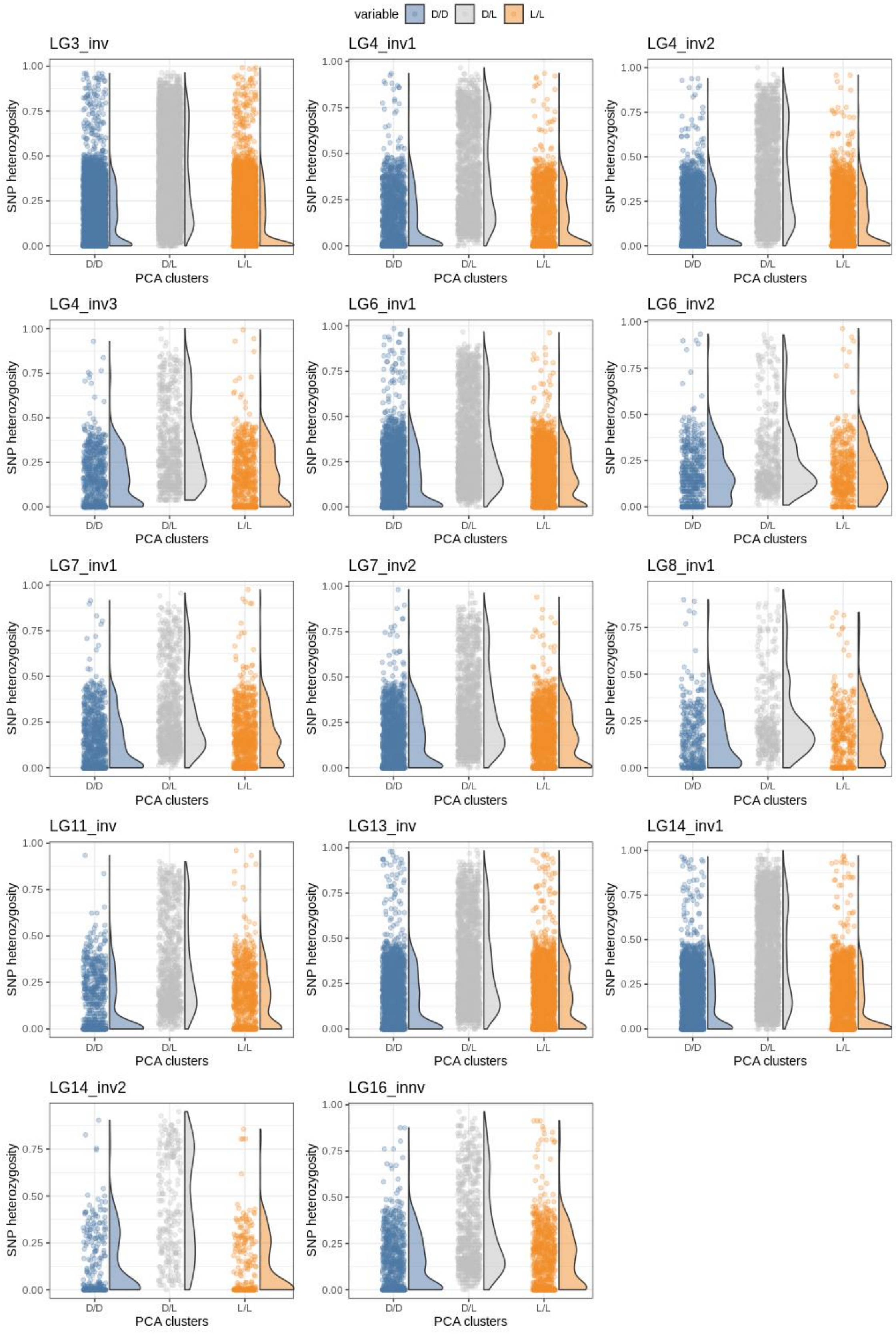

Figure S10: Average SNP heterozygosity per DAPC cluster. The cluster D/L always shows numerous SNPs with high heterozygosity (>0.7) despite the low coverage of the data (~5X).

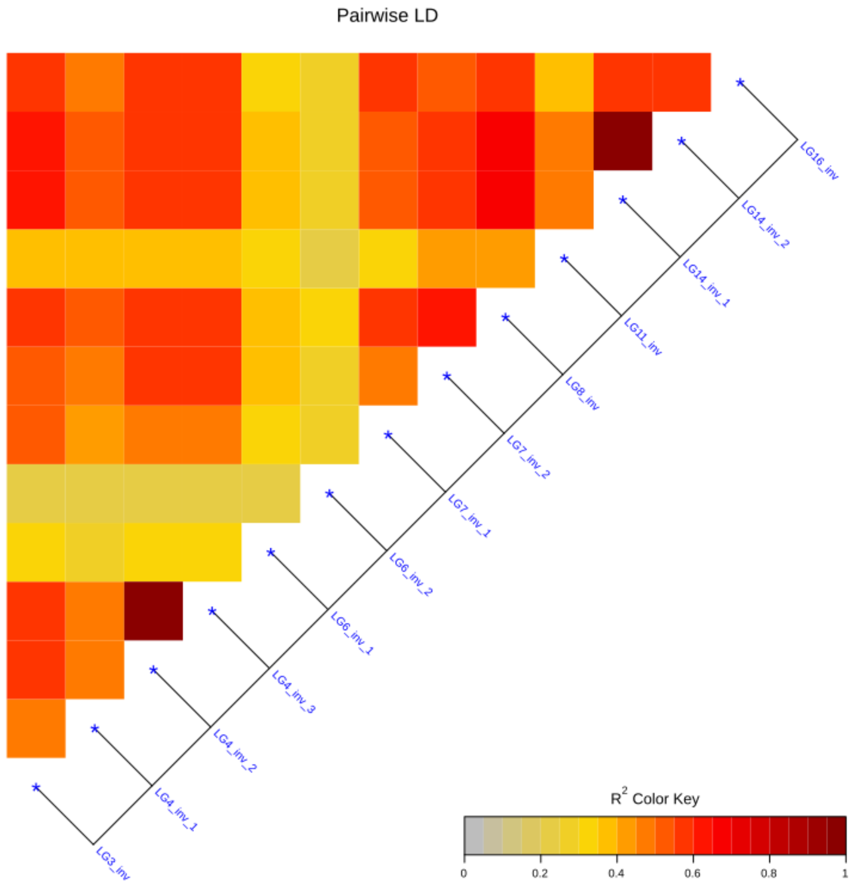

Figure S11: LD heatmap between putative rearrangements. LD was calculated using the R package LDheatmap based on local DAPC clusters as represented in Figure S7.

### Quantitative trait clines

#### Shell size

Table S3: Results of the shell length clines inferred by Bayesian inferences (with the Rstan package). For each fitted cline, the table shows the model selection statistics: the leave-one-out informatics criterion (looic), the differences in looic from the best model ($\Delta_{looic}$) and the loo weight ($W_{loo}$) followed by the estimated parameters of the clines, presented with their 95% posterior intervals. In order of appearance: the center of the cline (m), the width of the cline (m), the mean sizes (mm) in dwarf ecotype (left part of the transect, i.e., the most sheltered part) and in the large ecotype (right part of the transect, i.e., the exposed part), the standard deviations of size in the left (sl) and right (sr) of the cline and the elevation in variance in the center (hybrid zone, sh$^2$). The line(s) in bold highlight(s) the best fitting model(s) in each transect.

| Transect | cline | looic | $\Delta_{loo}$ | $W_{loo}$ | Estimates | centre | w | left | right | sl | sh | sr |
|---|---|---|---|---|---|---|---|---|---|---|---|---|
| South | uni | -238.088 | -13.056 | 0.000 | mean | 111.34 | 55.65 | 8.85 | 11.16 | 1.23 | 1.51 | 1.29 |
| | | | | | 2.50% | 88.84 | 1.62 | 8.44 | 10.42 | 0.80 | 1.15 | 0.24 |
| | | | | | 97.50% | 132.99 | 108.93 | 9.28 | 12.27 | 1.64 | 1.74 | 1.74 |
| | **bini** | **-225.032** | **0.000** | **0.809** | **mean** | **122.82** | **94.59** | **8.72** | **11.83** | **0.98** | **-** | **0.88** |
| | | | | | 2.50% | 107.36 | 65.06 | 8.46 | 11.35 | 0.80 | - | 0.61 |
| | | | | | 97.50% | 137.17 | 112.02 | 8.99 | 12.15 | 1.21 | - | 1.30 |
| | bimo | -232.176 | -7.144 | 0.088 | mean | 109.32 | 75.63 | 8.72 | 11.26 | 1.08 | - | 1.46 |
| | | | | | 2.50% | 94.94 | 33.36 | 8.43 | 10.71 | 0.84 | - | 0.93 |
| | | | | | 97.50% | 126.67 | 110.52 | 9.04 | 11.95 | 1.35 | - | 1.73 |
| | **trini** | **-225.780** | **-0.748** | **0.103** | **mean** | **123.60** | **98.70** | **8.65** | **11.94** | **0.95** | **1.53** | **0.63** |
| | | | | | 2.50% | 110.27 | 72.72 | 8.38 | 11.59 | 0.75 | 0.99 | 0.32 |
| | | | | | 97.50% | 135.62 | 112.46 | 8.93 | 12.26 | 1.20 | 1.74 | 1.15 |
| | trimo | -233.458 | -8.426 | 0.000 | mean | 117.15 | 88.84 | 8.75 | 11.60 | 1.04 | 1.60 | 0.99 |
| | | | | | 2.50% | 98.12 | 38.37 | 8.46 | 10.78 | 0.76 | 1.23 | 0.31 |
| | | | | | 97.50% | 132.76 | 112.28 | 9.08 | 12.15 | 1.37 | 1.74 | 1.72 |
| North | uni | -298.952 | -12.607 | 0.008 | mean | 62.80 | 60.36 | 8.10 | 11.51 | 1.07 | 1.34 | 1.36 |
| | | | | | 2.50% | 47.14 | 18.94 | 7.13 | 11.13 | 0.18 | 1.09 | 0.26 |
| | | | | | 97.50% | 74.40 | 110.92 | 8.85 | 11.94 | 1.64 | 1.63 | 1.82 |
| | **bini** | **-287.008** | **-0.663** | **0.424** | **mean** | **69.21** | **77.71** | **8.40** | **11.60** | **0.93** | **-** | **1.01** |
| | | | | | 2.50% | 57.29 | 49.53 | 8.12 | 11.32 | 0.73 | - | 0.83 |
| | | | | | 97.50% | 80.32 | 109.85 | 8.72 | 11.83 | 1.20 | - | 1.23 |
| | bimo | -293.316 | -6.971 | 0.000 | mean | 62.75 | 62.77 | 8.33 | 11.40 | 0.95 | - | 1.28 |
| | | | | | 2.50% | 52.38 | 27.20 | 7.94 | 11.13 | 0.69 | - | 1.10 |
| | | | | | 97.50% | 72.81 | 113.27 | 8.73 | 11.68 | 1.28 | - | 1.51 |
| | **trini** | **-286.345** | **0.000** | **0.568** | **mean** | **64.21** | **94.60** | **7.95** | **11.73** | **0.65** | **1.34** | **0.94** |
| | | | | | 2.50% | 52.74 | 64.03 | 7.59 | 11.46 | 0.40 | 0.80 | 0.77 |
| | | | | | 97.50% | 76.05 | 118.61 | 8.43 | 11.99 | 1.04 | 1.78 | 1.16 |
| | trimo | -293.703 | -7.358 | 0.000 | mean | 61.71 | 87.08 | 8.09 | 11.52 | 0.77 | 1.52 | 1.15 |
| | | | | | 2.50% | 49.23 | 38.72 | 7.53 | 11.18 | 0.41 | 0.88 | 0.81 |
| | | | | | 97.50% | 74.57 | 119.05 | 8.64 | 11.84 | 1.25 | 1.82 | 1.47 |

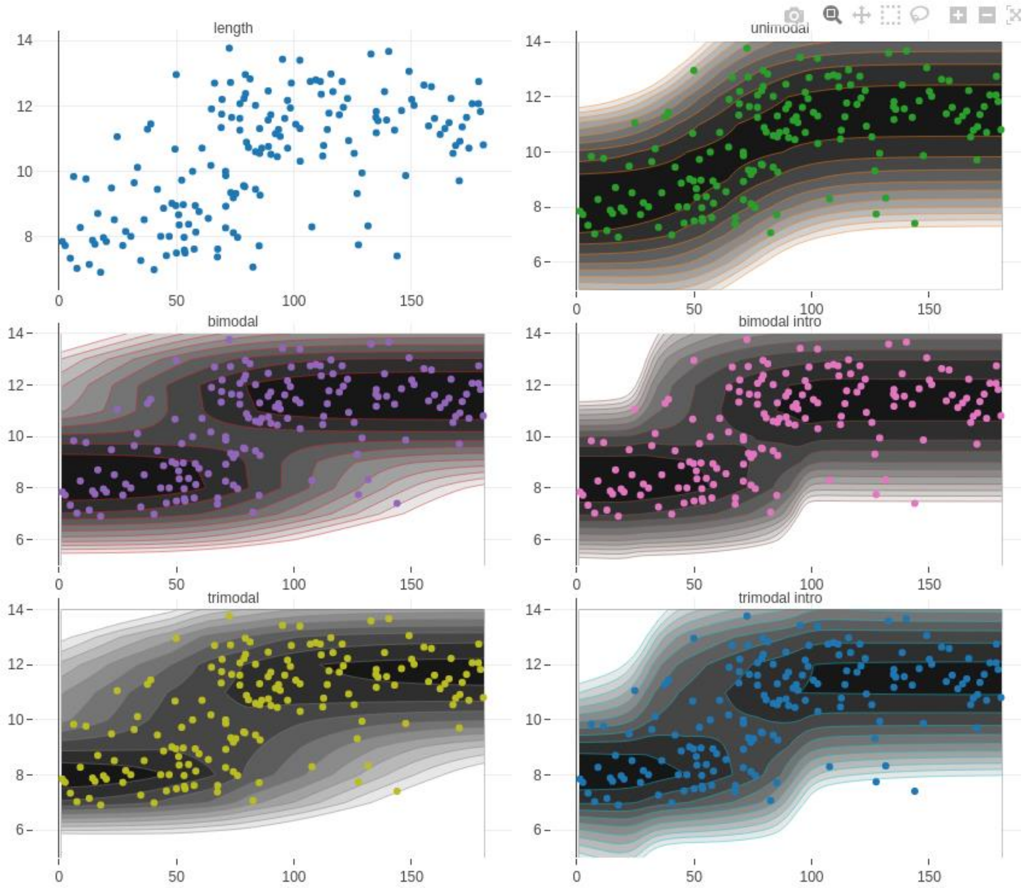

Figure S12: Multimodal clines for shell size along the northern transect. Each dot represents the size of a snail plotted against its position along the transect. The shade of grey represents the probability of observing a given size given the shape of the cline and the position on the transect. For this transect, the best fitting clines are the bimodal and the trimodal clines both without introgression (middle and bottom left).

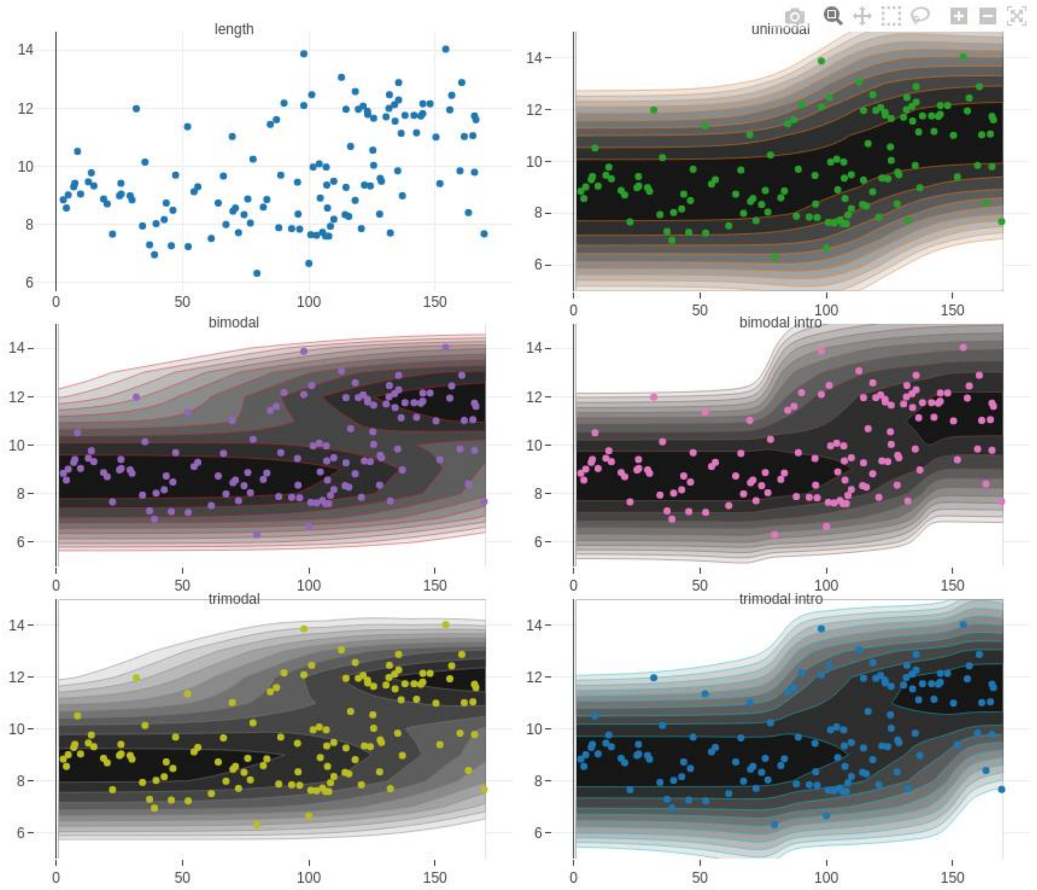

Figure S13: Multimodal clines for shell size along the southern transect. Each dot represents the size of a snail plotted against its position along the transect. The shade of grey represents the probability of observing a given size given the shape of the cline and the position on the transect. For this transect, the best fitting cline is a bimodal cline without introgression.

#### PC1 score

**Table S4: Results of the PC1 score clines** inferred by Bayesian inferences (with the Rstan package). For each fitted cline, the table shows the model selection statistics: the leave-one-out informatics criterion (looic), the differences in looic from the best model ($\Delta_{looic}$) and the loo weight ($W_{loo}$) followed by the estimated parameters of the clines, presented with their 95% posterior intervals. In order of appearance: the center of the cline (m), the width of the cline (m), the mean PC score in dwarf ecotype (left part of the transect, i.e., the most sheltered part) and in the large ecotype (right part of the transect, i.e., the exposed part), the standard deviations of PC score in the left (sl) and right (sr) of the cline and the elevation in variance in the center (hybrid zone, sh$^2$). The line(s) in bold highlight(s) the best fitting model(s) in each transect.

| transect | cline | looic | $\Delta_{looic}$ | $W_{loo}$ | Estimates | centre | w | left | right | sl | sh | sr |
|---|---|---|---|---|---|---|---|---|---|---|---|---|
| South | uni | -613,17 | -91,36 | 0,00 | mean | 105,91 | 66,74 | -58,50 | 49,64 | 13,26 | 48,87 | 5,51 |
| | | | | | 2.5% | 97,36 | 40,65 | -64,68 | 41,88 | 0,37 | 44,45 | 0,11 |
| | | | | | 97.5% | 114,63 | 90,01 | -48,91 | 55,43 | 30,12 | 50,37 | 20,27 |
| | **bini** | **-543,66** | **-21,86** | **0,18** | **mean** | **103,67** | **62,18** | **-57,51** | **39,47** | **7,96** | | **18,25** |
| | | | | | **2.5%** | **95,30** | **43,04** | **-59,38** | **34,71** | **6,61** | | **14,64** |
| | | | | | **97.5%** | **111,70** | **88,84** | **-55,58** | **44,02** | **9,73** | | **22,67** |
| | **bi** | **-541,70** | **-19,90** | **0,17** | **mean** | **92,30** | **92,15** | **-58,21** | **40,61** | **7,22** | | **18,27** |
| | | | | | **2.5%** | **90,34** | **64,46** | **-60,03** | **35,37** | **5,62** | | **14,57** |
| | | | | | **97.5%** | **94,90** | **111,84** | **-56,20** | **45,57** | **9,23** | | **23,04** |
| | **trini** | **-521,80** | **0,00** | **0,45** | **mean** | **112,36** | **89,98** | **-58,77** | **51,56** | **5,99** | **34,26** | **3,15** |
| | | | | | 2.5% | 103,45 | 69,62 | -60,43 | 50,49 | 4,59 | 25,78 | 2,42 |
| | | | | | 97.5% | 123,64 | 110,26 | -56,96 | 52,67 | 7,51 | 45,13 | 4,12 |
| | **tri** | **-526,96** | **-5,16** | **0,20** | **mean** | **106,62** | **99,52** | **-59,51** | **51,64** | **5,17** | **35,71** | **3,16** |
| | | | | | **2.5%** | **97,94** | **75,95** | **-61,89** | **50,49** | **2,64** | **27,22** | **2,42** |
| | | | | | **97.5%** | **114,20** | **112,48** | **-57,37** | **52,80** | **7,13** | **46,45** | **4,15** |
| North | uni | -763,65 | -66,49 | 0,00 | mean | 64,88 | 57,47 | -61,04 | 47,70 | 3,97 | 46,53 | 12,69 |
| | | | | | 2.5% | 60,91 | 47,32 | -65,90 | 43,57 | 0,14 | 43,78 | 9,40 |
| | | | | | 97.5% | 69,08 | 70,26 | -55,58 | 51,61 | 13,51 | 47,69 | 16,68 |
| | **bini** | **-749,00** | **-51,84** | **0,20** | **mean** | **65,81** | **44,97** | **-51,34** | **42,32** | **13,42** | | **17,71** |
| | | | | | 2.5% | 58,92 | 30,68 | -55,48 | 38,05 | 9,28 | | 13,45 |
| | | | | | 97.5% | 72,61 | 65,31 | -45,73 | 46,44 | 19,55 | | 21,92 |
| | bi | -734,25 | -37,09 | 0,07 | mean | 63,68 | 61,36 | -58,69 | 47,19 | 11,48 | | 15,99 |
| | | | | | 2.5% | 57,82 | 38,75 | -63,97 | 42,69 | 8,65 | | 13,07 |
| | | | | | 97.5% | 68,91 | 99,64 | -53,50 | 51,68 | 14,99 | | 19,65 |
| | trini | -709,27 | -12,10 | 0,00 | mean | 68,21 | 62,64 | -57,58 | 48,55 | 5,24 | 29,18 | 9,81 |
| | | | | | 2.5% | 61,01 | 45,37 | -59,27 | 45,56 | 3,94 | 20,37 | 3,42 |
| | | | | | 97.5% | 73,86 | 108,60 | -55,82 | 53,36 | 6,81 | 39,66 | 12,91 |
| | **tri** | **-697,16** | **0,00** | **0,72** | **mean** | **70,19** | **113,63** | **-58,36** | **53,82** | **4,48** | **30,61** | **3,12** |
| | | | | | 2.5% | 61,54 | 99,43 | -60,18 | 52,55 | 3,15 | 24,72 | 2,32 |
| | | | | | 97.5% | 76,92 | 120,08 | -56,44 | 54,96 | 6,18 | 37,42 | 4,27 |

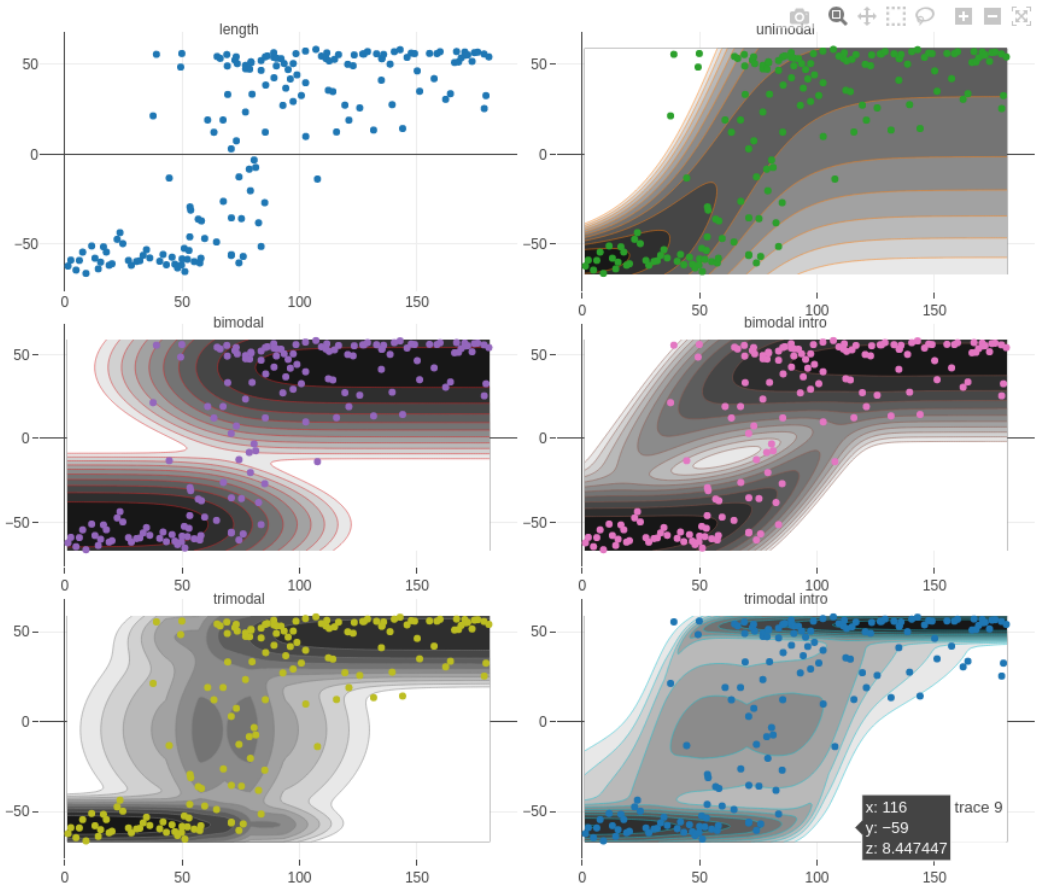

Figure S14: Multimodal cline for PC1 score along the northern transect. Each dot represents the PC score of ~~the~~ a snail~~s~~ in the PCA ~~analyses~~ analysis in Figure 1b plotted against its position along the transect. The shade of grey represents the probability of observing a given PC score given the shape of the cline and the position on the transect. For this transect, the best ~~fitted~~ fitting cline is the trimodal cline with introgression (bottom right), closely followed by the bimodal clines without introgression (in the middle left).

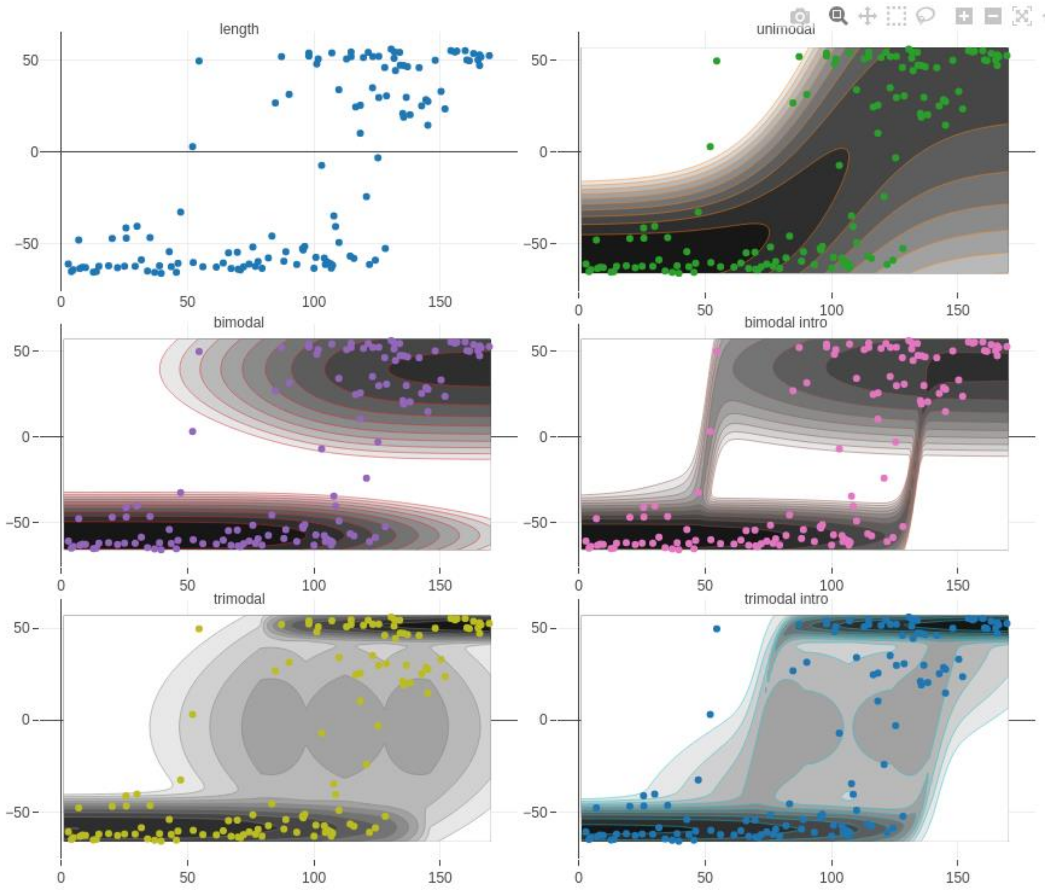

Figure S15: Multimodal cline for PC1 score along the southern transect. Each dot represents the PC score of ~~the~~ a snail~~s~~ in the PCA ~~analyses~~ analysis in Figure 1b plotted against its position along the transect. The shade of grey represents the probability of observing a given PC score given the shape of the cline and the position on the transect. For this transect, the best fitted cline is the trimodal cline without introgression (bottom left), closely followed by the trimodal cline with introgression (bottom right) and the two bimodal clines (with and without introgression, in the middle).

#### Hybrid index

Table S5: Results of the hybrid index clines inferred by Bayesian inferences (with Rstan package) for the southern and northern transect. We computed the hybrid index using four different datasets: the collinear outlier with $F_{ST}$ value above the 95% quantile of $F_{ST}$, the inversion karyotypes, all outliers above the 95% quantile and above the 99% quantile. For each fitted cline, inferred by Bayesian inferences (with the Rstan package). For each fitted cline, the table shows the model selection statistics: the leave-one-out informatics criterion (looic), the differences in looic from the best model ($\Delta_{looic}$) and the loo weight ($W_{loo}$) followed by the estimated parameters of the clines, presented with their 95% posterior intervals. In order of appearance: the center of the cline (m), the width of the cline (m), the mean hybrid index in dwarf ecotype (left part of the transect, i.e., the most sheltered part) and in the large ecotype (right part of the transect, i.e., the exposed part), the standard deviations in hybrid index the left (sl) and right (sr) of the cline and the elevation in variance in the center (hybrid zone, $sh^2$). The line(s) in bold highlight(s) the best fitting model(s) in each transect.

| transect | dataset | cline | loo | $\Delta_{loo}$ | $W_{loo}$ | Estimates | centre | w | left | right | sl | sh | sr |
|---|---|---|---|---|---|---|---|---|---|---|---|---|---|
| South | Colinear | uni | 79,73 | -42,51 | 0,07 | mean | 109,79 | 83,88 | 0,11 | 0,97 | 7,64 | -16,99 | 11,51 |
| | | | | | | 2,50% | 100,24 | 59,09 | 0,06 | 0,90 | 4,45 | -24,85 | 6,84 |
| | | | | | | 97,50% | 118,57 | 109,34 | 0,17 | 1,00 | 12,01 | -10,27 | 16,81 |
| | | **bini** | **118,85** | **-3,39** | **0,40** | **mean** | **104,50** | **60,65** | **0,10** | **0,87** | **17,16** | | **11,64** |
| | | | | | | 2,50% | 95,75 | 40,95 | 0,09 | 0,84 | 12,14 | | 7,16 |
| | | | | | | 97,50% | 112,57 | 87,89 | 0,12 | 0,90 | 19,90 | | 17,30 |
| | | bimo | 92,80 | -29,44 | 0,00 | mean | 105,60 | 97,65 | 0,12 | 0,88 | 15,96 | 7,69 | 0,43 |
| | | | | | | 2,50% | 102,77 | 50,63 | 0,09 | 0,85 | 6,29 | 4,03 | 0,22 |
| | | | | | | 97,50% | 109,11 | 112,68 | 0,15 | 0,91 | 19,92 | 19,18 | 0,50 |
| | | **trini** | **122,24** | **0,00** | **0,54** | **mean** | **104,45** | **63,55** | **0,10** | **0,89** | **18,24** | **6,14** | **16,24** |
| | | | | | | 2,50% | 96,26 | 44,84 | 0,08 | 0,87 | 14,51 | 0,89 | 10,93 |
| | | | | | | 97,50% | 112,53 | 88,30 | 0,11 | 0,91 | 19,93 | 18,67 | 19,83 |
| | | tri | 113,91 | -8,33 | 0,00 | mean | 105,35 | 86,56 | 0,10 | 0,89 | 18,19 | 1,71 | 17,08 |
| | | | | | | 2,50% | 100,30 | 55,43 | 0,09 | 0,86 | 14,20 | 0,78 | 10,83 |
| | | | | | | 97,50% | 110,55 | 111,57 | 0,12 | 0,91 | 19,94 | 3,18 | 19,89 |
| South | AllInv | uni | 79,73 | -42,51 | 0,07 | mean | 109,79 | 83,88 | 0,11 | 0,97 | 7,64 | -16,99 | 11,51 |
| | | | | | | 2,50% | 100,24 | 59,09 | 0,06 | 0,90 | 4,45 | -24,85 | 6,84 |
| | | | | | | 97,50% | 118,57 | 109,34 | 0,17 | 1,00 | 12,01 | -10,27 | 16,81 |
| | | **bini** | **118,85** | **-3,39** | **0,40** | **mean** | **104,50** | **60,65** | **0,10** | **0,87** | **17,16** | | **11,64** |
| | | | | | | 2,50% | 95,75 | 40,95 | 0,09 | 0,84 | 12,14 | | 7,16 |
| | | | | | | 97,50% | 112,57 | 87,89 | 0,12 | 0,90 | 19,90 | | 17,30 |
| | | bimo | 92,80 | -29,44 | 0,00 | mean | 105,60 | 97,65 | 0,12 | 0,88 | 15,96 | 7,69 | 0,43 |
| | | | | | | 2,50% | 102,77 | 50,63 | 0,09 | 0,85 | 6,29 | 4,03 | 0,22 |
| | | | | | | 97,50% | 109,11 | 112,68 | 0,15 | 0,91 | 19,92 | 19,18 | 0,50 |
| | | **trini** | **122,24** | **0,00** | **0,54** | **mean** | **104,45** | **63,55** | **0,10** | **0,89** | **18,24** | **6,14** | **16,24** |
| | | | | | | 2,50% | 96,26 | 44,84 | 0,08 | 0,87 | 14,51 | 0,89 | 10,93 |
| | | | | | | 97,50% | 112,53 | 88,30 | 0,11 | 0,91 | 19,93 | 18,67 | 19,83 |
| | | tri | 113,91 | -8,33 | 0,00 | mean | 105,35 | 86,56 | 0,10 | 0,89 | 18,19 | 1,71 | 17,08 |
| | | | | | | 2,50% | 100,30 | 55,43 | 0,09 | 0,86 | 14,20 | 0,78 | 10,83 |
| | | | | | | 97,50% | 110,55 | 111,57 | 0,12 | 0,91 | 19,94 | 3,18 | 19,89 |
| South | Q99 | uni | 188,90 | -41,12 | 0,02 | mean | 112,54 | 76,69 | 0,10 | 0,97 | 2,56 | -5,75 | 4,36 |

| | | | | | | | | | | | | | |
|---|---|---|---|---|---|---|---|---|---|---|---|---|---|
| | | | | | | 2,50% | 102,37 | 50,25 | 0,04 | 0,89 | 1,39 | -8,63 | 2,52 |
| | | | | | | 97,50% | 121,98 | 106,67 | 0,19 | 1,00 | 4,13 | -3,22 | 6,40 |
| | | **bini** | **229,37** | **-0,65** | **0,55** | **mean** | **104,73** | **62,43** | **0,03** | **0,89** | **17,94** | | **4,19** |
| | | | | | | 2,50% | 96,14 | 43,32 | 0,03 | 0,84 | 13,53 | | 2,32 |
| | | | | | | 97,50% | 112,75 | 87,13 | 0,04 | 0,93 | 19,91 | | 6,64 |
| | | bimo | 184,47 | -45,55 | 0,00 | mean | 106,77 | 84,64 | 0,07 | 0,86 | 10,69 | | 2,56 |
| | | | | | | 2,50% | 102,32 | 48,74 | 0,03 | 0,78 | 1,94 | | 1,29 |
| | | | | | | 97,50% | 111,56 | 111,76 | 0,14 | 0,92 | 19,89 | | 5,79 |
| | | **trini** | **230,02** | **0,00** | **0,43** | **mean** | **108,25** | **65,44** | **0,03** | **0,94** | **18,64** | **2,37** | **8,83** |
| | | | | | | 2,50% | 99,60 | 45,21 | 0,02 | 0,90 | 15,43 | 0,55 | 4,05 |
| | | | | | | 97,50% | 117,23 | 91,99 | 0,04 | 0,97 | 19,97 | 8,92 | 18,52 |
| | | tri | 206,43 | -23,59 | 0,00 | mean | 106,87 | 76,99 | 0,03 | 0,94 | 16,62 | 0,74 | 9,74 |
| | | | | | | 2,50% | 101,77 | 45,63 | 0,02 | 0,90 | 8,64 | 0,41 | 4,03 |
| | | | | | | 97,50% | 113,31 | 110,25 | 0,05 | 0,97 | 19,92 | 1,29 | 18,45 |
| South | Q95 | **uni** | **299,31** | **-19,92** | **0,12** | **mean** | **108,83** | **77,03** | **0,11** | **0,96** | **1,70** | **-3,39** | **2,57** |
| | | | | | | 2,50% | 97,07 | 47,94 | 0,03 | 0,88 | 0,91 | -5,19 | 1,43 |
| | | | | | | 97,50% | 119,05 | 107,27 | 0,20 | 1,00 | 2,75 | -1,70 | 3,85 |
| | | **bini** | **319,22** | **0,00** | **0,88** | **mean** | **102,85** | **61,48** | **0,03** | **0,89** | **12,63** | | **2,30** |
| | | | | | | 2,50% | 93,93 | 42,06 | 0,02 | 0,82 | 6,02 | | 1,19 |
| | | | | | | 97,50% | 111,33 | 86,60 | 0,05 | 0,93 | 18,83 | | 3,83 |
| | | bimo | 303,50 | -15,72 | 0,00 | mean | 101,95 | 78,27 | 0,03 | 0,82 | 11,88 | | 1,08 |
| | | | | | | 2,50% | 84,75 | 46,48 | 0,02 | 0,71 | 5,05 | | 0,60 |
| | | | | | | 97,50% | 109,78 | 109,74 | 0,05 | 0,89 | 19,16 | | 1,76 |
| | | trini | 318,97 | -0,25 | 0,00 | mean | 105,44 | 66,31 | 0,02 | 0,93 | 14,84 | 2,31 | 4,07 |
| | | | | | | 2,50% | 96,89 | 45,01 | 0,02 | 0,88 | 8,20 | 0,41 | 1,83 |
| | | | | | | 97,50% | 114,06 | 95,10 | 0,04 | 0,97 | 19,76 | 10,04 | 8,20 |
| | | tri | 307,49 | -11,73 | 0,00 | mean | 106,41 | 69,22 | 0,04 | 0,93 | 10,13 | 0,56 | 3,75 |
| | | | | | | 2,50% | 99,60 | 42,54 | 0,02 | 0,86 | 2,71 | 0,31 | 1,54 |
| | | | | | | 97,50% | 112,52 | 107,31 | 0,08 | 0,97 | 18,76 | 0,97 | 8,03 |
| North | Colinear | uni | 116,46 | -37,89 | 0,00 | mean | 65,18 | 65,15 | 0,09 | 0,90 | 9,51 | -18,79 | 11,68 |
| | | | | | | 2,50% | 56,93 | 33,75 | 0,01 | 0,87 | 3,83 | -29,17 | 8,21 |
| | | | | | | 97,50% | 74,02 | 92,09 | 0,20 | 0,93 | 16,83 | -10,34 | 15,85 |
| | | bini | 139,24 | -15,11 | 0,00 | mean | 67,92 | 48,91 | 0,15 | 0,88 | 9,88 | | 15,21 |
| | | | | | | 2,50% | 61,15 | 33,53 | 0,12 | 0,85 | 5,89 | | 9,32 |
| | | | | | | 97,50% | 74,28 | 68,92 | 0,19 | 0,90 | 16,03 | | 19,72 |
| | | bimo | 128,58 | -25,77 | 0,00 | mean | 58,55 | 91,08 | 0,11 | 0,89 | 14,07 | | 14,61 |
| | | | | | | 2,50% | 54,58 | 47,10 | 0,08 | 0,86 | 4,58 | | 9,16 |
| | | | | | | 97,50% | 63,97 | 119,95 | 0,14 | 0,91 | 19,58 | | 19,72 |
| | | **trini** | **154,35** | **0,00** | **0,97** | **mean** | **67,03** | **46,13** | **0,11** | **0,89** | **17,13** | **4,61** | **17,85** |
| | | | | | | 2,50% | 60,12 | 33,33 | 0,09 | 0,88 | 12,03 | 1,44 | 14,12 |
| | | | | | | 97,50% | 72,24 | 61,59 | 0,13 | 0,91 | 19,90 | 15,65 | 19,91 |
| | | tri | 142,51 | -11,84 | 0,03 | mean | 65,35 | 73,73 | 0,12 | 0,89 | 17,77 | 2,16 | 18,18 |
| | | | | | | 2,50% | 56,68 | 44,17 | 0,09 | 0,87 | 13,10 | 1,04 | 14,45 |
| | | | | | | 97,50% | 71,54 | 116,48 | 0,14 | 0,90 | 19,93 | 4,25 | 19,94 |
| North | AllInv | uni | 116,46 | -37,89 | 0,00 | mean | 65,18 | 65,15 | 0,09 | 0,90 | 9,51 | -18,79 | 11,68 |

| | | | | | | | | | | | | | |
|---|---|---|---|---|---|---|---|---|---|---|---|---|---|
| | | | | | | 2,50% | 56,93 | 33,75 | 0,01 | 0,87 | 3,83 | -29,17 | 8,21 |
| | | | | | | 97,50% | 74,02 | 92,09 | 0,20 | 0,93 | 16,83 | -10,34 | 15,85 |
| | | bini | 139,24 | -15,11 | 0,00 | mean | 67,92 | 48,91 | 0,15 | 0,88 | 9,88 | | 15,21 |
| | | | | | | 2,50% | 61,15 | 33,53 | 0,12 | 0,85 | 5,89 | | 9,32 |
| | | | | | | 97,50% | 74,28 | 68,92 | 0,19 | 0,90 | 16,03 | | 19,72 |
| | | bimo | 128,58 | -25,77 | 0,00 | mean | 58,55 | 91,08 | 0,11 | 0,89 | 14,07 | | 14,61 |
| | | | | | | 2,50% | 54,58 | 47,10 | 0,08 | 0,86 | 4,58 | | 9,16 |
| | | | | | | 97,50% | 63,97 | 119,95 | 0,14 | 0,91 | 19,58 | | 19,72 |
| | | **trini** | **154,35** | **0,00** | **0,97** | **mean** | **67,03** | **46,13** | **0,11** | **0,89** | **17,13** | **4,61** | **17,85** |
| | | | | | | 2,50% | 60,12 | 33,33 | 0,09 | 0,88 | 12,03 | 1,44 | 14,12 |
| | | | | | | 97,50% | 72,24 | 61,59 | 0,13 | 0,91 | 19,90 | 15,65 | 19,91 |
| | | tri | 142,51 | -11,84 | 0,03 | mean | 65,35 | 73,73 | 0,12 | 0,89 | 17,77 | 2,16 | 18,18 |
| | | | | | | 2,50% | 56,68 | 44,17 | 0,09 | 0,87 | 13,10 | 1,04 | 14,45 |
| | | | | | | 97,50% | 71,54 | 116,48 | 0,14 | 0,90 | 19,93 | 4,25 | 19,94 |
| North | Q99 | uni | 236,46 | -24,65 | 0,00 | mean | 64,98 | 74,49 | 0,06 | 0,93 | 3,36 | -6,40 | 4,27 |
| | | | | | | 2,50% | 57,26 | 47,43 | 0,00 | 0,90 | 1,47 | -9,55 | 2,91 |
| | | | | | | 97,50% | 74,36 | 100,14 | 0,17 | 0,96 | 5,28 | -3,45 | 5,80 |
| | | bini | 243,61 | -17,50 | 0,00 | mean | 58,82 | 43,29 | 0,04 | 0,84 | 15,66 | | 2,36 |
| | | | | | | 2,50% | 51,79 | 30,17 | 0,02 | 0,79 | 3,52 | | 1,59 |
| | | | | | | 97,50% | 67,54 | 62,14 | 0,11 | 0,90 | 19,86 | | 3,96 |
| | | bimo | 245,61 | -15,50 | 0,07 | mean | 59,39 | 51,69 | 0,03 | 0,85 | 16,04 | | 2,43 |
| | | | | | | 2,50% | 54,14 | 33,57 | 0,02 | 0,80 | 5,98 | | 1,69 |
| | | | | | | 97,50% | 64,39 | 80,30 | 0,07 | 0,89 | 19,92 | | 3,51 |
| | | **trini** | **261,11** | **0,00** | **0,61** | **mean** | **68,51** | **62,45** | **0,03** | **0,95** | **17,75** | **7,91** | **10,90** |
| | | | | | | 2,50% | 62,02 | 44,58 | 0,02 | 0,92 | 12,74 | 3,48 | 4,88 |
| | | | | | | 97,50% | 74,28 | 93,93 | 0,04 | 0,97 | 19,93 | 15,15 | 19,56 |
| | | **tri** | **259,20** | **-1,91** | **0,32** | **mean** | **61,93** | **62,23** | **0,03** | **0,92** | **17,94** | **9,94** | **5,05** |
| | | | | | | 2,50% | 58,33 | 41,71 | 0,02 | 0,88 | 13,47 | 3,34 | 3,06 |
| | | | | | | 97,50% | 66,96 | 91,07 | 0,04 | 0,94 | 19,93 | 18,58 | 7,61 |
| North | Q95 | uni | 383,93 | -15,84 | 0,00 | mean | 61,70 | 68,83 | 0,06 | 0,93 | 2,20 | -3,81 | 2,55 |
| | | | | | | 2,50% | 53,53 | 42,40 | 0,00 | 0,89 | 0,94 | -5,99 | 1,69 |
| | | | | | | 97,50% | 72,38 | 92,84 | 0,21 | 0,96 | 3,57 | -1,85 | 3,59 |
| | | bini | 387,26 | -12,52 | 0,00 | mean | 63,10 | 39,53 | 0,07 | 0,89 | 5,99 | | 2,12 |
| | | | | | | 2,50% | 53,67 | 26,26 | 0,02 | 0,83 | 1,93 | | 1,22 |
| | | | | | | 97,50% | 71,93 | 57,88 | 0,14 | 0,93 | 16,28 | | 3,48 |
| | | bimo | 387,70 | -12,07 | 0,00 | mean | 60,19 | 55,31 | 0,06 | 0,89 | 6,51 | | 2,09 |
| | | | | | | 2,50% | 54,85 | 31,66 | 0,02 | 0,84 | 2,59 | | 1,29 |
| | | | | | | 97,50% | 65,66 | 99,64 | 0,11 | 0,93 | 15,88 | | 3,27 |
| | | **trini** | **399,67** | **-0,10** | **0,46** | **mean** | **66,07** | **50,81** | **0,03** | **0,94** | **15,00** | **5,06** | **4,22** |
| | | | | | | 2,50% | 59,66 | 36,00 | 0,02 | 0,90 | 7,38 | 1,79 | 2,18 |
| | | | | | | 97,50% | 71,14 | 66,84 | 0,04 | 0,96 | 19,76 | 9,72 | 6,85 |
| | | **tri** | **399,77** | **0,00** | **0,54** | **mean** | **61,37** | **60,25** | **0,02** | **0,93** | **15,66** | **5,19** | **3,29** |
| | | | | | | 2,50% | 57,65 | 37,80 | 0,02 | 0,89 | 8,43 | 2,09 | 1,83 |
| | | | | | | 97,50% | 66,36 | 92,61 | 0,04 | 0,96 | 19,83 | 10,12 | 5,19 |

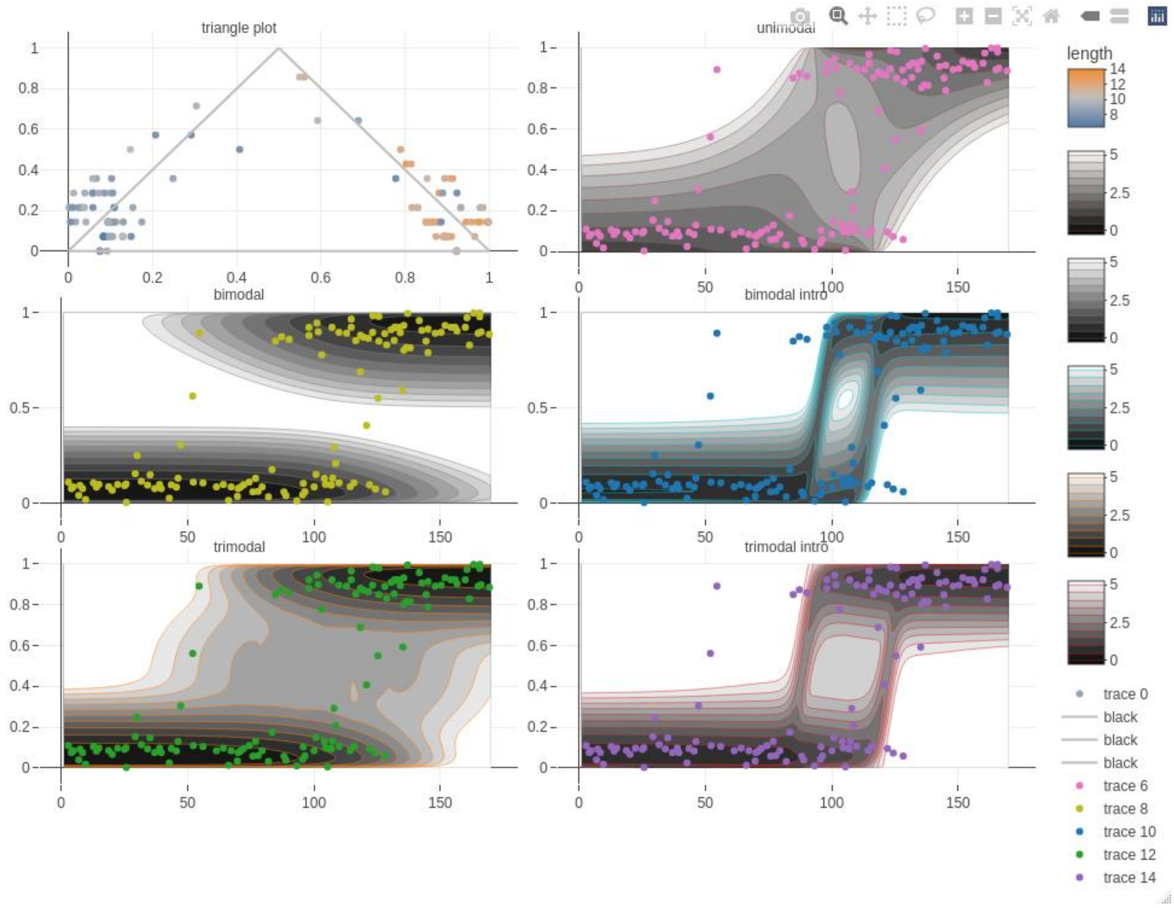

Figure S16: Multimodal cline for the hybrid index along the southern transect. Each dot represents the hybrid index computed using the inversion karyotypes of the snails, which was plotted with estimates of inter-specific heterozygosity in the triangle plots (top left), and against its position along the transect for the clines analyses. The shade of grey represents the probability of observing a given hybrid index given the shape of the cline and the position on the transect. For this transect, the best fitted cline is the trimodal cline without introgression (bottom left), closely followed by the bimodal clines without introgression (in the middle left). Similar clines were obtained when the hybrid index was calculated based on other datasets, thus, the shape of the clines ~~are~~ is not showed here.

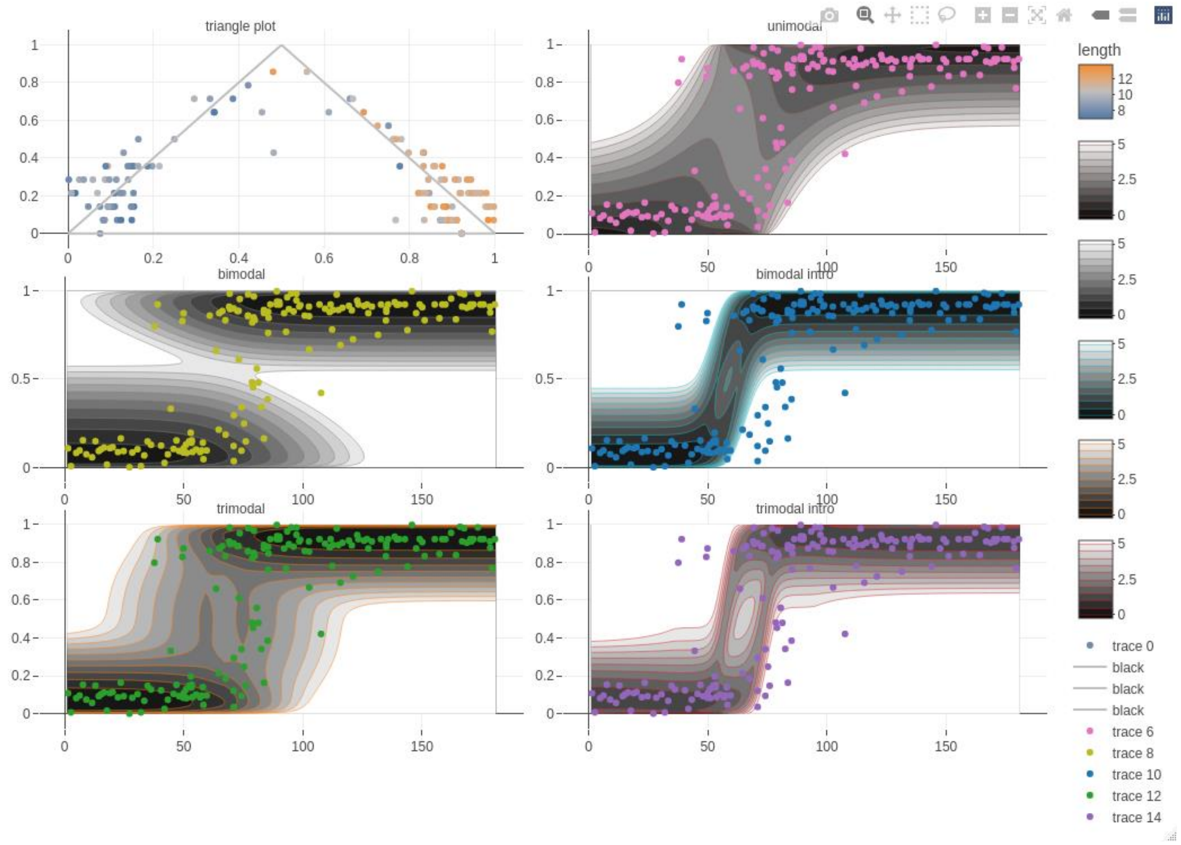

Figure S17: Multimodal cline for the hybrid index along the northern transect**.** Each dot represents the hybrid index computed using the inversion karyotypes of the snails, which was plotted with estimates of inter-specific heterozygosity in the triangle plots (top left), and against its position along the transect for the clines analyses. The shade of grey represents the probability of observing a given hybrid index given the shape of the cline and the position on the transect. For this transect, the best fitted cline is the trimodal cline without introgression (bottom left). Similar clines were obtained when the hybrid index was calculated based on other datasets, thus, the shape of the clines is~~are~~ not showed here.

### Genetic clines

#### Arrangement clines

Table S6: Results of the arrangement cline fitting for the inversion karyotypes inferred from clustering patterns in the PCA analyses shown in Figure S7. The cline fits were performed with the R package Rstan and included an estimate of the cline center in meters (center), cline width in meters (width), the frequency in the sheltered part of the shore where the dwarf ecotype is located (Dwarf) and in the exposed area where the large ecotype is located (large). The cline fit also included an estimate of $F_{IS}$ assumed to peak at the cline centre and estimated at 5 points, following the formulation in Cfit. $F_{IS}$ values significantly different from 0 at the centre (with 95% posterior interval excluding 0) are highlighted by a *. The following columns show the goodness of the cline fit assessed by the percentage of deviance explained using generalize linear regression model with binomial error (Dev. Explain) and the estimates of the selection coefficient (s) using a per generation dispersal of 8.27m (see below).

| Transect | Data | center | width | dwarf | large | $F_{IS}$ | Dev. Exp. | s |
|---|---|---|---|---|---|---|---|---|
| North | LG3_inv | 68,05 | 57,75 | 0,99 | 0,08 | **0,52*** | 46,05 | 0,051 |
| | LG4_inv1 | 59,6 | 75,56 | 0,97 | 0,03 | **0,53*** | 43,48 | 0,032 |
| | LG4_inv2 | 60,99 | 55,99 | 0,97 | 0,06 | **0,61*** | 46,38 | 0,054 |
| | LG4_inv3 | 60,99 | 55,49 | 0,97 | 0,06 | **0,61*** | 46,38 | 0,055 |
| | LG6_inv1 | 57 | 88,71 | 0,88 | 0,1 | 0,22 | 25,16 | 0,016 |
| | LG6_inv2 | 72,24 | 29,06 | 0,59 | 0,13 | -0,43 | 23,82 | 0,051 |
| | LG7_inv1 | 62,15 | 65,47 | 0,97 | 0,11 | **0,59*** | 39,1 | 0,035 |
| | LG7_inv2 | 61,27 | 64,45 | 0,99 | 0,08 | **0,48*** | 43,94 | 0,041 |
| | LG8_inv | 65,83 | 60,97 | 0,99 | 0,05 | **0,7*** | 47,45 | 0,049 |
| | LG11_inv | 63,55 | 59,93 | 0,74 | 0,04 | 0,4 | 36,61 | 0,028 |
| | LG13_inv | 64,83 | 74,18 | 0,92 | 0,04 | **0,4*** | 41,05 | 0,029 |
| | LG14_inv1 | 69,62 | 78,89 | 0,99 | 0,01 | **0,63*** | 46,73 | 0,032 |
| | LG14_inv2 | 69,56 | 79,1 | 0,99 | 0,01 | **0,62*** | 46,73 | 0,031 |
| | LG16_innv | 64,78 | 42,78 | 0,99 | 0,09 | **0,5*** | 51,63 | 0,091 |
| South | LG3_inv | 108,79 | 72,93 | 1 | 0,06 | **0,75*** | 45 | 0,034 |
| | LG4_inv1 | 104,15 | 72,75 | 0,9 | 0,05 | **0,73*** | 36,25 | 0,028 |
| | LG4_inv2 | 99,05 | 88,87 | 0,99 | 0,01 | **0,66*** | 41,24 | 0,025 |
| | LG4_inv3 | 98,77 | 89,23 | 0,99 | 0,01 | **0,66*** | 41,24 | 0,025 |
| | LG6_inv1 | 107,25 | 75,9 | 0,76 | 0,02 | 0,42 | 30,65 | 0,020 |
| | LG6_inv2 | 104,37 | 66,74 | 0,76 | 0,08 | 0,43 | 26,9 | 0,021 |
| | LG7_inv1 | 99,82 | 81,7 | 0,99 | 0,11 | **0,69*** | 39,01 | 0,024 |
| | LG7_inv2 | 105,01 | 89,96 | 0,98 | 0,01 | **0,84*** | 39,01 | 0,024 |
| | LG8_inv | 103,92 | 79,16 | 1 | 0,01 | **0,89*** | 44,41 | 0,032 |
| | LG11_inv | 108,55 | 78,37 | 0,75 | 0,01 | 0,5 | 27,97 | 0,018 |
| | LG13_inv | 99,27 | 91,89 | 0,98 | 0,02 | **0,73*** | 40,14 | 0,022 |
| | LG14_inv1 | 101,48 | 85,89 | 0,97 | 0,01 | **0,86*** | 42,88 | 0,026 |
| | LG14_inv2 | 101,12 | 86,62 | 0,97 | 0,01 | **0,86*** | 42,88 | 0,025 |
| | LG16_innv | 103,74 | 73,14 | 0,99 | 0,03 | **0,73*** | 47,29 | 0,035 |

#### SNPs allelic clines

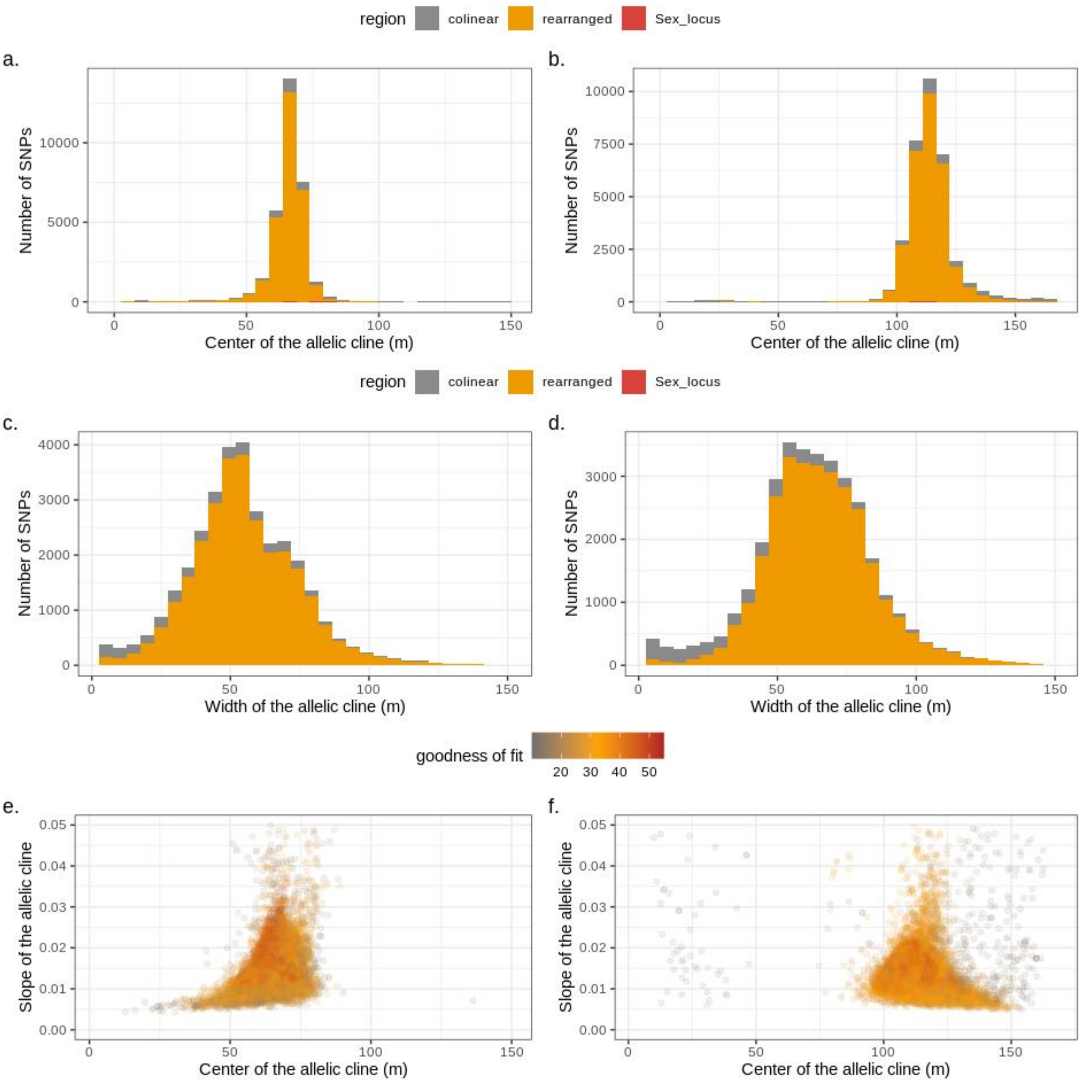

Figure S18: Allele frequency cline fits for individual SNPs for the northern (a,-c,-e) and southern (b,-d,-f) transects. a. and b. show the distributions of cline centers, c. and d. the distributions of cline widths. In a. to d., colors distinguish SNPs from collinear (grey) and rearranged (orange). e. and f. show the slopes of the cline (differences of allele frequency / width) against the center of the cline. In e. and f. each dot corresponds to a SNP that is colored by the percentage of deviance explained by the cline fit.

#### Estimates of dispersal

An allele frequency cline allows estimation (from the cline width) of the selection coefficient (*s*) required to maintain the rate of change of allele frequency observed across a transect, assuming a balance between selection and gene flow. However, prior knowledge about dispersal, specifically an estimate of the standard deviation of parent – offspring distances in the direction of the transect (σ), is required to estimate *s* from the cline width. This is an important constraint (Barton and Hewitt 1985, Barton and Gale 1993). The marine snail *L. fabalis* has a non-overlapping generation time of one year, which means that parents generally die before their offspring mature. In our data, we found 12 pairs of closely-related individuals (Figure S19). Because *L. fabalis* has non-overlapping generations, and as we sampled mature individuals, the closely related individuals match the signature expected from full-sibs and half-sibs. This provided a rare opportunity to obtain an estimate of dispersal from an undisturbed natural population. We used the distance between siblings to infer the parent-offspring distance using the following procedure.

Assuming offspring dispersal follows a normal distribution $N(0, \sigma^2)$ in the direction of the transect, then the distances between pairs of offspring from the same parent is also a normal distribution of $N(0, \sigma_o^2)$, where $\sigma_o^2 = 2\ \sigma^2$. Furthermore, the absolute distance along the transect is a half-normal distribution $N(\bar{d},\sigma_d{}^2)$, where $\bar{d}$ is the mean absolute distance between sibling samples, and $\sigma_d{}^2$ is the variance in absolute distance. The two parameters $\bar{d}$ and $\sigma_d{}^2$ can be related to the normal distribution $N(0, 2\sigma^2)$:

$$\sigma_d^2 = 2\sigma^2\left(1 - \frac{2}{\pi}\right) \text{ and } \bar{d} = \sqrt{2}\ \sigma\ \sqrt{2/\pi}$$

Thus, we can obtain an estimate of σ from the distances between sibling or half-sib pairs using the following formula:

$$\sigma = \frac{\bar{d}}{\sqrt{2}\ \sqrt{2/\pi}} = \frac{\bar{d}}{2/\sqrt{\pi}}$$

The average distance between closely related individuals ($\bar{d}$) was 16.99m (min = 2.96m, max = 101.23m). However, one full-sib pair was a clear outlier (indicated by the arrow in Fig. S19). This suggests a long tail to the dispersal distribution that violates our assumption of normality, and may be important for colonization of new habitats as well as influencing patterns of LD in the hybrid zone but has less influence on cline width (Barton and Gale 1993). After dropping this outlier, the mean distance between sib pairs was 9.33m (max=28.54m). Using $\bar{d} = 9.33m$, we estimated that the standard deviation of parent-offspring distances, σ was 8.27m with a standard error of 2.01m, estimated by jackknife resampling over sibling pairs.

The number of sibling pairs observed also provides an estimate of the short-term effective size of the population, because the proportion of individuals in the current generation that is expected to share one or both parents depends on the number of parents contributing offspring. We obtained an estimate of the population size in each transect by simulating 100000 samples of the same size as our experimental samples from populations of different sizes (N, ranging from 500 to 8000 individuals), with equal sex ratio, random mating and Poisson distributed offspring number using a custom R script and a linear model relating 1/N to the numbers of full-sib pairs, half-sib pairs and half-sib individuals.

For the observed numbers of siblings, we obtained estimates of population size of 5138 snails (CI: 5036-5243) in the northern transect and 5059 snails (CI: 4961-5167) in the southern transect.

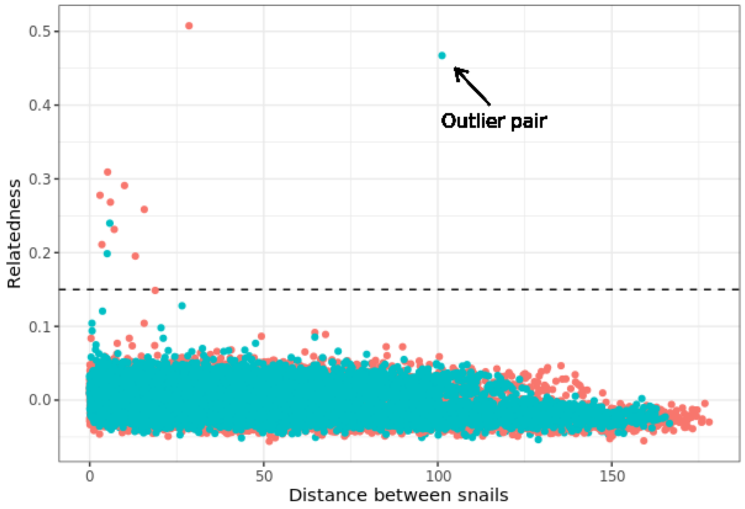

Figure S19: Pairwise relatedness against the transect distance between pairs of individuals from the northern shore (red) and southern shore (blue). Relatedness values above the dashed line is expected from pair of siblings. 12 pairs showed a relatedness value above this expected value (10 half sib pairs and two full sib pairs).

#### Association between barrier loci

Linkage disequilibrium between loci in hybrid zones is generated primarily by dispersal. It generates attraction between clines causing them to become coupled and further enhancing the LD due to the increased overlap (Barton et al. Bierne et al.). This disequilibrium (D) reaches a maximum in the center of a hybrid zone, and the strength of D depends on the width of the hybrid zone (w), the per generation dispersal (σ), and the recombination between different loci (r). In the absence of epistatic interaction between barrier loci or assortative mating, and when measured after dispersal, the expected D ($D_{exp}$) can be calculated as follows (Barton and Gale, 1993):

$$a.\ D_{exp} = (\sigma^2(1+r))/(r * w^2).$$

Assuming free recombination (r=0.5), and using our estimate of σ = 8.27 (±2.01, Figure S19), *w* from the unimodal inversion clines of 83.81 in the southern shore (±13.70, Table S5) and 71.35 in the northern shore (±12.51, Table S5), we obtain a $D_{exp}$ of 0.029 and 0.040 for the southern and northern shore, respectively. The confidence intervals of $D_{exp}$ (Table S7) were then calculated by randomly sampling 1000 combinations from the confidence intervals of *w* and σ, and assuming that both *w* and σ are normally distributed around their mean estimates.

$D_{obs}$ at any specific position along a hybrid zone can be extracted from the hybrid index using (Barton and Gale, 1993):

$$b.\ D_{obs} = \frac{(var(z)-\frac{1}{n}(\bar{z}(1-\bar{z})-var(p)))}{\frac{1}{2}(1-\frac{1}{n})}.$$

Here, z is the hybrid index, n is the number of loci (in this case, n=9 which corresponds to the 9 nearly fixed different inversions, after removing the two inversions from LG6 and the inversion on LG11 showing intermediate frequency in the dwarf ecotype), and var(p) is the variance in arrangement frequency across the 9 inversions.

$D_{obs}$ is expected to reach a maximum in the center of the hybrid zone, where $\bar{z} = 0.5$, and the mean frequency of each karyotype is also 0.5 (resulting in the expected heterozygosity of 0.5). To calculate the variance in hybrid index (var(z)) and the variance in karyotype frequency (var(p)) at the centre of the hybrid zone, we first split our data into bins containing 10 consecutive snails along the transects. We then calculated, for each bin, the mean frequency of the inversion arrangement ($\bar{p}$), the variance in arrangement frequencies across the 9 inversions (var(p)), and the variance in hybrid index (var(z)). $\bar{p}$ was then used to generate expected heterozygosity ($2\bar{p}\,(1-\bar{p})$), and this was used to predict the var(p) and var(z) at the zone centre (expected heterozygosity = 0.5) using linear regression. The results are summarised in table S7. Overall, the observed disequilibrium between the main barrier loci is much higher than the expected value of D estimated from the dispersal and the cline widths. These results point toward some mechanism increasing the association between barrier loci located on different chromosomes, and suggests the presence of assortative mating or epistatic interaction between barrier loci, or both.

Table S7: Summary table of the association between fixed inversion. In order of appearance, the table shows the hybrid zone studied, the variance in arrangement frequencies (var(p)) at the centre of the hybrid zone for the 9 nearly-fixed inversions and estimated from a linear regression of this variance on expected heterozygosity, the standard error of the var(p) estimates, the variance in hybrid index (var(z)) at the centre of the hybrid zone, also estimated from a linear regression on expected, the standard error of var(z) estimates, the observed disequilibrium ($D_{obs}$) calculated from the equation b., the standard error of $D_{obs}$, the expected disequilibrium ($D_{exp}$), the upper and the lower 2.5% confidence interval of $D_{exp.}$

| HZ | var(p) | se var(p) | var(z) | se var(z) | $D_{obs}$ | se Dobs | $D_{exp}$ | Upper | Lower |
|---|---|---|---|---|---|---|---|---|---|
| North | 0.007 | 0.001 | 0.129 | 0.014 | 0.259 | 0.043 | 0.04 | 0.128 | 0.01 |
| South | 0.005 | 0.002 | 0.183 | 0.017 | 0.382 | 0.049 | 0.029 | 0.086 | 0.008 |

#### Demographic inference

Table S8 Summary table of the demographic inferences. In order of appearance, the table shows the dataset used, the model tested, 4 columns related to the model fit and comparison: the likelihood (likeli.), the Akaike criterion (AIC), the differences in AIC (ΔAIC) and the AIC weight (wAIC), and 15 columns with the parameters estimated from the model (all expressed relative to theta): the ancestral population mutation rate (theta), the ancestral population size after size variation (Naf), the effective population sizes of the large ecotype (Ne1) and the dwarf ecotype (Ne2), the time at which the variation in ancestral effective population size happened (Tae), the time of split (Ts), the time of secondary contact (Tsc), the effective number of migrants from large to dwarf (M1) and from dwarf to large (M2), the proportion by which M1 or M2 is reduced inside genomic islands (Mr1, and Mr2), the proportion of the genome affected by heterogeneous gene flow (P), the Hill-Robertson effect approximated here by the fraction by which the effective population sizes N1 and N2 are reduced (hrf), and the proportion of the genome with reduced effective population size (Q). Parameters not estimated in a given model were set to NA. Best fitted model is highlighted in bold.

| Population | Model | likeli. | AIC | ΔAIC | wAIC | Theta | Naf | Ne1 | Ne2 | Tae | Ts | Tsc | M1 | M2 | Mr1 | Mr2 | P | hrf | Q |
|---|---|---|---|---|---|---|---|---|---|---|---|---|---|---|---|---|---|---|---|
| overall | SI | -23696 | 47397 | 42207 | 0 | 9851 | NA | 11.89 | 0.57 | NA | 0.07 | NA | NA | NA | NA | NA | NA | NA | NA |
| | IM | -14315 | 28641 | 23451 | 0 | 4270 | NA | 98.45 | 0.01 | NA | 1.16 | NA | 8.53 | 189.63 | NA | NA | NA | NA | NA |
| | AM | -16494 | 33000 | 27810 | 0 | 4007 | NA | 0.21 | 8.36 | NA | 1.03 | 0 | 16.08 | 14.27 | NA | NA | NA | NA | NA |
| | SC | -12253 | 24518 | 19328 | 0 | 5457 | NA | 7.25 | 0.07 | NA | 0.61 | 0.02 | 4.78 | 56.26 | NA | NA | NA | NA | NA |
| | SI_ae | -19182 | 38373 | 33183 | 0 | 4401 | 5.9 | 99.7 | 0.5 | 1.01 | 0.06 | NA | NA | NA | NA | NA | NA | NA | NA |
| | IM_ae | -14304 | 28622 | 23432 | 0 | 3293 | 3.68 | 81.01 | 0.01 | 1.21 | 0.93 | NA | 6.54 | 191.68 | NA | NA | NA | NA | NA |
| | AM_ae | -14560 | 29135 | 23945 | 0 | 929 | 11.37 | 55.58 | 0.48 | 7.95 | 3.05 | 0 | 0.33 | 5.27 | NA | NA | NA | NA | NA |
| | SC_ae | -11756 | 23529 | 18338 | 0 | 4557 | 5.74 | 99.42 | 0.09 | 0.96 | 0.16 | 0.02 | 3.03 | 46.94 | NA | NA | NA | NA | NA |
| | SI_2N | -9562 | 19134 | 13944 | 0 | 9034 | NA | 99.5 | 53.43 | NA | 0.27 | NA | NA | NA | NA | NA | NA | 0.01 | 0.19 |
| | IM_2N | -3799 | 7612 | 2422 | 0 | 7643 | NA | 4.48 | 2.94 | NA | 0.75 | NA | 3.85 | 4.1 | NA | NA | NA | 0.03 | 0.5 |
| | AM_2N | -3967 | 7950 | 2759 | 0 | 7267 | NA | 4.07 | 3.9 | NA | 0.78 | 0 | 4.19 | 3.94 | NA | NA | NA | 0.03 | 0.5 |
| | SC_2N | -2996 | 6009 | 819 | 0 | 5517 | NA | 6.32 | 2.43 | NA | 1.16 | 0.14 | 4.38 | 9.67 | NA | NA | NA | 0.02 | 0.5 |
| | SI_ae_2N | -4668 | 9350 | 4160 | 0 | 9406 | 0.27 | 9.57 | 0.07 | 1.75 | 0.05 | NA | NA | NA | NA | NA | NA | 0.98 | 0.23 |
| | IM_ae_2N | -3034 | 6087 | 896 | 0 | 8198 | 0.22 | 99.82 | 0.02 | 1.11 | 0.36 | NA | 1.42 | 12.97 | NA | NA | NA | 0.07 | 0.31 |
| | AM_ae_2N | -3349 | 6717 | 1527 | 0 | 7555 | 0.22 | 99.77 | 0.03 | 0.87 | 0.4 | 0 | 1.54 | 12.74 | NA | NA | NA | 0.07 | 0.36 |
| | SC_ae_2N | -3001 | 6022 | 831 | 0 | 5453 | 15.79 | 5.3 | 3.5 | 0.01 | 1.21 | 0.15 | 6.92 | 5.38 | NA | NA | NA | 0.01 | 0.5 |
| | IM_2M | -3417 | 6851 | 1660 | 0 | 4369 | NA | 99.55 | 0.13 | NA | 0.99 | NA | 31.28 | 195.94 | 0.01 | 0.01 | 0.31 | NA | NA |
| | AM_2M | -3562 | 7142 | 1952 | 0 | 4497 | NA | 99.79 | 0.25 | NA | 0.93 | 0 | 38.02 | 159.09 | 0.04 | 0.01 | 0.34 | NA | NA |
| | SC_2M | -2748 | 5515 | 324 | 0 | 4428 | NA | 5.44 | 0.37 | NA | 0.86 | 0.01 | 21.87 | 154.76 | 0.05 | 0.16 | 0.26 | NA | NA |
| | IM_ae_2M | -3395 | 6811 | 1621 | 0 | 4024 | 1.63 | 99.07 | 0.18 | 0.7 | 0.89 | NA | 33.6 | 188.98 | 0.01 | 0.01 | 0.32 | NA | NA |
| | AM_ae_2M | -3645 | 7313 | 2123 | 0 | 3360 | 1.4 | 97.39 | 0.28 | 1.11 | 1.17 | 0 | 30.16 | 155.85 | 0.01 | 0 | 0.32 | NA | NA |
| | SC_ae_2M | -2742 | 5505 | 315 | 0 | 4519 | 0.01 | 5.24 | 0.38 | 0 | 0.91 | 0.01 | 21.99 | 154.43 | 0.05 | 0.16 | 0.26 | NA | NA |

| | | | | | | | | | | | | | | | | | | | |
|---|---|---|---|---|---|---|---|---|---|---|---|---|---|---|---|---|---|---|---|
| | IM_2M_2N | -3089 | 6199 | 1008 | 0 | 8271 | NA | 0.08 | 0.03 | NA | 1.29 | NA | 4.81 | 12.48 | 0.96 | 1 | 0.13 | 78.96 | 0.32 |
| | AM_2M_2N | -3997 | 8017 | 2827 | 0 | 1923 | NA | 0.29 | 93.88 | NA | 1.1 | 0 | 76.01 | 16.02 | 0.01 | 0.01 | 0.48 | 0.83 | 0.05 |
| | **SC_2M_2N** | **-2584** | **5190** | **0** | **1** | **2649** | **NA** | **5.88** | **0.44** | **NA** | **0.94** | **0.01** | **17.41** | **130.72** | **0.06** | **0.16** | **0.33** | **0** | **0.49** |
| | IM_ae_2M_2N | -3132 | 6288 | 1097 | 0 | 18620 | 0.25 | 99.51 | 0.04 | 3.37 | 0.45 | NA | 171.53 | 59.51 | 0.13 | 0.01 | 0.24 | 0.19 | 0.04 |
| | AM_ae_2M_2N | -4236 | 8499 | 3308 | 0 | 2120 | 19.21 | 0.5 | 76.6 | 0.41 | 0.7 | 0 | 66.06 | 14.08 | 0.01 | 0.01 | 0.41 | 0.11 | 0.11 |
| | SC_ae_2M_2N | -2695 | 5416 | 226 | 0 | 3705 | 0 | 3.92 | 0.51 | 0 | 0.91 | 0.02 | 70.3 | 78.45 | 0.09 | 0.05 | 0.32 | 0 | 0.4 |
| colinear | SI | -7273 | 14552 | 11483 | 0 | 7096 | NA | 9.63 | 45.23 | NA | 0.17 | NA | NA | NA | NA | NA | NA | NA | NA |
| | IM | -3134 | 6279 | 3209 | 0 | 3026 | NA | 1.76 | 97.96 | NA | 1.18 | NA | 5.4 | 5.23 | NA | NA | NA | NA | NA |
| | AM | -3360 | 6731 | 3662 | 0 | 3069 | NA | 0.6 | 6.89 | NA | 0.99 | 0 | 36.76 | 0.09 | NA | NA | NA | NA | NA |
| | SC | -2895 | 5802 | 2733 | 0 | 3138 | NA | 6.17 | 1.09 | NA | 1.09 | 0.05 | 15.01 | 15.76 | NA | NA | NA | NA | NA |
| | SI_ae | -3368 | 6747 | 3677 | 0 | 3061 | 6.57 | 4.23 | 99.25 | 0.98 | 0.1 | NA | NA | NA | NA | NA | NA | NA | NA |
| | IM_ae | -3076 | 6165 | 3096 | 0 | 131673 | 0.02 | 0.06 | 48.92 | 0.27 | 0.03 | NA | 129.54 | 191.56 | NA | NA | NA | NA | NA |
| | AM_ae | -3108 | 6233 | 3163 | 0 | 2719 | 4.93 | 1.49 | 97.91 | 1.04 | 0.48 | 0 | 9.33 | 3.93 | NA | NA | NA | NA | NA |
| | SC_ae | -2893 | 5803 | 2733 | 0 | 1743 | 3.18 | 11.88 | 1.71 | 2.14 | 1.65 | 0.09 | 8.37 | 10.36 | NA | NA | NA | NA | NA |
| | SI_2N | -5649 | 11308 | 8238 | 0 | 7046 | NA | 35.25 | 97.74 | NA | 0.27 | NA | NA | NA | NA | NA | NA | 0.02 | 0.05 |
| | IM_2N | -1836 | 3685 | 616 | 0 | 4908 | NA | 3.82 | 3.4 | NA | 1.09 | NA | 9.86 | 7.52 | NA | NA | NA | 0.02 | 0.43 |
| | AM_2N | -1850 | 3717 | 647 | 0 | 4709 | NA | 3.81 | 3.53 | NA | 1.11 | 0 | 10.1 | 7.42 | NA | NA | NA | 0.02 | 0.42 |
| | SC_2N | -1691 | 3398 | 328 | 0 | 3251 | NA | 4.12 | 3.05 | NA | 1.11 | 0.07 | 16.64 | 8.04 | NA | NA | NA | 0.01 | 0.14 |
| | SI_ae_2N | -1831 | 3676 | 606 | 0 | 3032 | 10.77 | 98.13 | 0.06 | 1.08 | 0.04 | NA | NA | NA | NA | NA | NA | 0.07 | 0.91 |
| | IM_ae_2N | -1560 | 3138 | 68 | 0 | 3689 | 5.34 | 3.65 | 99.38 | 1.32 | 0.53 | NA | 5.27 | 4.36 | NA | NA | NA | 0.03 | 0.37 |
| | AM_ae_2N | -1596 | 3211 | 142 | 0 | 3950 | 5.57 | 3.03 | 99.7 | 1.01 | 0.45 | 0 | 6.64 | 5.62 | NA | NA | NA | 0.03 | 0.35 |
| | **SC_ae_2N** | **-1525** | **3070** | **0** | **0.998** | **3716** | **5.7** | **7.55** | **99.86** | **1.16** | **0.33** | **0.04** | **8.78** | **7.1** | **NA** | **NA** | **NA** | **0.02** | **0.3** |
| | IM_2M | -1768 | 3552 | 483 | 0 | 3257 | NA | 0.78 | 97.89 | NA | 0.94 | NA | 28.2 | 13.44 | 0.01 | 0.02 | 0.07 | NA | NA |
| | AM_2M | -1773 | 3563 | 493 | 0 | 3166 | NA | 0.8 | 99.15 | NA | 0.96 | 0 | 28.89 | 13.43 | 0.01 | 0.01 | 0.06 | NA | NA |
| | SC_2M | -1569 | 3156 | 86 | 0 | 6064 | NA | 36.07 | 0.45 | NA | 1.21 | 0.02 | 110.01 | 15.23 | 0.47 | 0.04 | 0.05 | NA | NA |
| | IM_ae_2M | -1690 | 3400 | 330 | 0 | 43038 | 0.07 | 0.09 | 76.11 | 5.52 | 0.08 | NA | 197.56 | 141.35 | 0.02 | 0.01 | 0.07 | NA | NA |
| | AM_ae_2M | -1719 | 3460 | 390 | 0 | 2630 | 5.18 | 0.4 | 99.82 | 1.2 | 0.34 | 0 | 84.52 | 12.11 | 0.01 | 0.01 | 0.05 | NA | NA |
| | SC_ae_2M | -1568 | 3158 | 89 | 0 | 5766 | 8.32 | 99.56 | 0.35 | 0.78 | 0.86 | 0.02 | 137.99 | 18.79 | 0.5 | 0.03 | 0.05 | NA | NA |
| | IM_2M_2N | -1664 | 3348 | 279 | 0 | 1003 | NA | 99.86 | 0.36 | NA | 7.3 | NA | 15.11 | 16.57 | 0.05 | 0.01 | 0.08 | 33.42 | 0.27 |
| | AM_2M_2N | -1802 | 3626 | 557 | 0 | 1394 | NA | 8.92 | 0.53 | NA | 1.46 | 0 | 12.61 | 114.29 | 0.01 | 0.03 | 0.15 | 0.13 | 0.47 |
| | SC_2M_2N | -1557 | 3137 | 67 | 0 | 3501 | NA | 99.07 | 0.41 | NA | 1.21 | 0.02 | 145.53 | 19.32 | 0.41 | 0.03 | 0.07 | 0.07 | 0.28 |

| | | | | | | | | | | | | | | | | | | | |
|---|---|---|---|---|---|---|---|---|---|---|---|---|---|---|---|---|---|---|---|
| | IM_ae_2M_2N | -1621 | 3266 | 197 | 0 | 12834 | 0.14 | 0.54 | 64.02 | 0.63 | 0.18 | NA | 18.68 | 43.17 | 0 | 0.63 | 0.05 | 0.03 | 0.5 |
| | AM_ae_2M_2N | -1653 | 3333 | 263 | 0 | 1411 | 0.52 | 94.22 | 0.37 | 0 | 6.65 | 0 | 18.65 | 3.56 | 0.21 | 0.01 | 0.06 | 0.32 | 0.25 |
| | SC_ae_2M_2N | -1528 | 3082 | 12 | 0.002 | 2457 | 1.22 | 34.81 | 0.4 | 2.69 | 0.77 | 0.04 | 65.57 | 11.81 | 0.38 | 0.03 | 0.09 | 0.96 | 0.11 |
| rearranged | SI | -6327 | 12661 | 8984 | 0 | 2615 | NA | 0.28 | 0.21 | NA | 0.06 | NA | NA | NA | NA | NA | NA | NA | NA |
| | IM | -3711 | 7431 | 3755 | 0 | 451 | NA | 3.19 | 1.81 | NA | 9.63 | NA | 0.16 | 0.32 | NA | NA | NA | NA | NA |
| | AM | -5132 | 10277 | 6600 | 0 | 1650 | NA | 0.08 | 81.31 | NA | 0.56 | 0 | 9.56 | 1.02 | NA | NA | NA | NA | NA |
| | SC | -2625 | 5262 | 1586 | 0 | 1528 | NA | 0.99 | 0.61 | NA | 0.8 | 0.04 | 2.61 | 3.52 | NA | NA | NA | NA | NA |
| | SI_ae | -6199 | 12407 | 8731 | 0 | 3055 | 0.01 | 0.31 | 0.25 | 0 | 0.07 | NA | NA | NA | NA | NA | NA | NA | NA |
| | IM_ae | -3405 | 6823 | 3147 | 0 | 76503 | 0.02 | 99.5 | 0 | 9.9 | 0.03 | NA | 24.37 | 198.62 | NA | NA | NA | NA | NA |
| | AM_ae | -3722 | 7461 | 3784 | 0 | 465 | 0.02 | 3.12 | 1.78 | 2.4 | 9.96 | 0 | 0.17 | 0.32 | NA | NA | NA | NA | NA |
| | SC_ae | -2561 | 5138 | 1462 | 0 | 6783 | 0 | 0.23 | 0.14 | 0 | 0.17 | 0.01 | 12.73 | 17.49 | NA | NA | NA | NA | NA |
| | SI_2N | -4528 | 9066 | 5390 | 0 | 2420 | NA | 8.28 | 6.14 | NA | 0.11 | NA | NA | NA | NA | NA | NA | 0.04 | 0.67 |
| | IM_2N | -2890 | 5795 | 2119 | 0 | 2124 | NA | 2.08 | 1.09 | NA | 0.33 | NA | 1.14 | 2.48 | NA | NA | NA | 0.12 | 0.49 |
| | AM_2N | -3075 | 6165 | 2489 | 0 | 1776 | NA | 2.28 | 1.26 | NA | 0.5 | 0 | 0.79 | 1.39 | NA | NA | NA | 0.17 | 0.5 |
| | SC_2N | -1872 | 3761 | 84 | 0 | 1656 | NA | 2.58 | 1.58 | NA | 0.67 | 0.03 | 3.95 | 4.64 | NA | NA | NA | 0.08 | 0.5 |
| | SI_ae_2N | -3973 | 7959 | 4283 | 0 | 4277 | 6.24 | 2.83 | 99.59 | 0.2 | 0.07 | NA | NA | NA | NA | NA | NA | 0.06 | 0.86 |
| | IM_ae_2N | -2011 | 4039 | 363 | 0 | 30113 | 0 | 0.11 | 0.13 | 0.01 | 0.03 | NA | 48.95 | 41.79 | NA | NA | NA | 0.08 | 0.45 |
| | AM_ae_2N | -2843 | 5706 | 2029 | 0 | 5490 | 0.22 | 97.92 | 0.07 | 2.75 | 0.21 | 0 | 0.8 | 4.46 | NA | NA | NA | 0.26 | 0.01 |
| | **SC_ae_2N** | **-1828** | **3676** | **0** | **1** | **3357** | **0.01** | **1.19** | **0.95** | **0.02** | **0.33** | **0.03** | **5.95** | **5.51** | **NA** | **NA** | **NA** | **0.09** | **0.48** |
| | IM_2M | -2838 | 5693 | 2016 | 0 | 17346 | NA | 0.09 | 0.06 | NA | 0.78 | NA | 199.65 | 33.81 | 0.18 | 0.02 | 0.42 | NA | NA |
| | AM_2M | -2884 | 5785 | 2109 | 0 | 2085 | NA | 0.8 | 0.53 | NA | 8.89 | 0 | 47.9 | 3.06 | 0.01 | 0.01 | 0.45 | NA | NA |
| | SC_2M | -1879 | 3776 | 100 | 0 | 700 | NA | 2.7 | 1.57 | NA | 4.64 | 0.1 | 37.89 | 1.37 | 0.7 | 0.02 | 0.49 | NA | NA |
| | IM_ae_2M | -2838 | 5696 | 2019 | 0 | 16551 | 12.81 | 0.09 | 0.06 | 5.94 | 1.15 | NA | 185.04 | 31.37 | 0.18 | 0.02 | 0.41 | NA | NA |
| | AM_ae_2M | -2873 | 5767 | 2091 | 0 | 1768 | 99 | 0.85 | 0.56 | 0.02 | 7.52 | 0 | 24.91 | 4.81 | 0.13 | 0.02 | 0.49 | NA | NA |
| | SC_ae_2M | -1996 | 4013 | 337 | 0 | 314 | 68.59 | 4.06 | 3.74 | 0.03 | 9.95 | 0.11 | 6.43 | 2.94 | 0.26 | 0.11 | 0.5 | NA | NA |
| | IM_2M_2N | -2151 | 4322 | 646 | 0 | 415 | NA | 0.94 | 0.58 | NA | 9.03 | NA | 0.38 | 0.63 | 0.95 | 0.98 | 0 | 52.35 | 0.09 |
| | AM_2M_2N | -3165 | 6351 | 2675 | 0 | 3267 | NA | 0.27 | 0.23 | NA | 1.05 | 0 | 31.39 | 2.19 | 0.82 | 0.03 | 0.49 | 0 | 0.35 |
| | SC_2M_2N | -2149 | 4319 | 643 | 0 | 895 | NA | 1.14 | 0.98 | NA | 0.51 | 0.01 | 12.41 | 12.34 | 0 | 0 | 0 | 0.07 | 0.49 |
| | IM_ae_2M_2N | -2776 | 5576 | 1900 | 0 | 1297 | 0.5 | 88.92 | 0.1 | 2.82 | 0.35 | NA | 0.24 | 3.56 | 0.93 | 0.23 | 0.19 | 0.35 | 0.06 |
| | AM_ae_2M_2N | -2917 | 5860 | 2183 | 0 | 472 | 5.01 | 1.49 | 1.21 | 0.77 | 9.97 | 0 | 0.25 | 19.04 | 0.13 | 0.08 | 0.06 | 0.01 | 0.02 |
| | SC_ae_2M_2N | -2163 | 4352 | 676 | 0 | 822 | 0.15 | 1.43 | 0.78 | 0.01 | 0.75 | 0.03 | 3.74 | 4.33 | 0.02 | 0.01 | 0 | 0.1 | 0.48 |

Table S9: Summary table of the transformation of each parameter estimated from the best demographic scenario in each dataset. In order of appearance, the reference population size (Nref) estimated from theta, the ancestral population size after size variation (Na), the effective population sizes of the large ecotype (Ne1) and the dwarf ecotype (Ne2), the time of split (Ts), the time of secondary contact (Tsc), the effective number of migrants from large to dwarf (M1) and from dwarf to large (M2), the per generation migration rate from large to dwarf (m1) and from dwarf to large (m2), and the reduced migration rate from large to dwarf (mr1) and dwarf to large (mr2).

| data | model | Nref | Na | N1 | N2 | Ts | Tsc | M1 | M2 | m1 | m2 | mr1 | mr2 |
|---|---|---|---|---|---|---|---|---|---|---|---|---|---|
| overall | SC_2M_2N | 81268 | -- | 477859 | 35758 | 152784 | 1625 | 51.18 | 28.75 | 0.00031 | 0.00017 | 0.00002 | 0.00003 |
| colinear | SC_ae_2N | 114002 | 649817 | 860722 | 11384338 | 75241 | 9120 | 33.14 | 354.50 | 0.00014 | 0.00155 | -- | -- |
| rearranged | SC_ae_2N | 102989 | 1029 | 122557 | 97839 | 67972 | 6179 | 3.540 | 2.61 | 0.00002 | 0.00001 | -- | -- |
